# CXCL-CXCR2 signaling drives cancer-endothelium interactions in SCLC metastatic seeding

**DOI:** 10.64898/2026.04.15.716394

**Authors:** Zhang Yang, Andy Xu, Nicholas Hughes, Chien-Wei Peng, Adrienne Visani, Susrutha Puthanmadhom Narayanan, Xue Wen Ng, Isabella Guppy, Chris Roberts, Yaoxuan You, Monte M. Winslow, David W. Piston, Jon Park, Zhonglin Lyu, Feng Chen, Li Ding, Rui Tang

## Abstract

Small cell lung cancer (SCLC) is a highly metastatic malignancy that frequently colonizes the liver, yet the signaling mechanisms that govern early metastatic seeding remain incompletely defined. During metastasis, disseminated cancer cells encounter endothelial cells (ECs) at the vascular-tissue interface, positioning cancer-endothelium crosstalk as a potential determinant of metastatic success. Here, we identify an SCLC-induced endothelial CXCL chemokine program that engages cancer-cell CXCR2-RAC1 signaling during metastatic seeding. Single-cell and bulk transcriptomic profiling showed that SCLC exposure induces CXCL chemokines in liver sinusoidal endothelial cells and other endothelial models. CXCL stimulation enhanced SCLC migration, trans-endothelial migration, F-actin remodeling, and tumor-endothelial contact. Using Metastasis-Originated Barcode Sequencing (MOBA-seq) coupled to genetic perturbation, we show that CXCR2 promotes metastatic seeding across multiple organs, with prominent effects in the liver. Transcriptomic, protein-level, cell-cell interaction, biosensor, and genetic analyses further support a model in which CXCR2 signaling promotes RAC1-dependent cytoskeletal remodeling. Pharmacologic CXCR2 blockade reduced SCLC migration, SCLC-endothelial interactions, liver metastatic seeding, and prolonged survival following intravenous SCLC transplantation. Together, these findings define a reciprocal cancer-endothelial signaling axis that links endothelial CXCL induction to tumor-intrinsic CXCR2-RAC1 activity and identify CXCR2 as a potential vulnerability for limiting metastatic establishment in SCLC.

## Introduction

Small cell lung cancer (SCLC) accounts for ∼15% of lung cancer cases in the United States and causes more than 200,000 deaths worldwide each year^1, 2^. At diagnosis, more than two-thirds of patients present with distant metastases, with the liver among the most frequent sites of dissemination^3, 4^. Genomic studies have revealed recurrent, near-universal inactivation of TP53 and RB1, along with additional mutational signatures during SCLC initiation and progression^5, 6^. However, the molecular mechanisms and genetic dependencies that drive organ-specific metastatic colonization, including in the liver, remain poorly understood. Patients with extensive-stage SCLC (ES-SCLC) are primarily treated with etoposide and platinum-based chemotherapy^7, 8^. Effective treatments for metastatic SCLC remain limited, resulting in a median survival of less than 7 months^1^. Defining signaling pathways that promote metastatic seeding and outgrowth may uncover actionable vulnerabilities for earlier intervention and improved treatment of extensive-stage disease.

Metastasis is a multistep process requiring cancer cell intravasation, survival during circulation, extravasation and seeding at distant sites before clonal expansion into clinically overt disease^9, 10^. Endothelial cells (ECs) line the inner surface of the vascular system and therefore constitute the first interface between disseminated cancer cells and organ-specific tumor microenvironments (TME)^11–14^. ECs play complex and context-dependent roles during the metastatic cascade. They can form an antimetastatic niche by acting as a physical barrier to extravasation and by secreting factors that restrain tumor cell proliferation^14, 15^. Conversely, ECs can support established metastatic lesions through angiogenesis and remodel inflammatory programs influencing immune surveillance and metastatic outgrowth^15, 16^. Notably, pan-cancer single-cell analyses have revealed substantial heterogeneity in EC states across tumor types, suggesting that cancer cells actively remodel endothelial compartment to construct permissive TMEs^17, 18^. Despite these advances, functional roles of ECs and reciprocal signaling mechanisms governing cancer-endothelium communication in the metastatic cascade remain poorly understood.

Chemokine signaling is a central regulator of immune-endothelial interactions during inflammation and immune surveillance. In the context of cancer metastasis, circulating tumor cells can co-opt these pathways to promote organ-specific homing and extravasation, strengthen cancer-endothelium interactions and stimulate angiogenesis^19, 20^. Among these pathways, the CXCR2 receptor and its ligands, CXCL chemokines, are key drivers of neutrophil recruitment in shaping the tumor immune microenvironment^21–23^. However, most work has focused on CXCR2-mediated immune cell trafficking, with tumor cells primarily viewed as sources of CXCL chemokines rather than as CXCR2-responsive cells. Emerging evidence suggests CXCR2 is highly expressed in neuroendocrine tumor cells and may promote a phenotypic switch towards a more invasive state^24^. Despite this, the precise role of tumor-intrinsic CXCR2 signaling in metastatic SCLC remains poorly defined. Given the availability of high-affinity CXCR2 inhibitors in clinical trials and the association of CXCR2 activity with worse patient outcomes, defining the role of CXCR2 signaling in metastatic progression could provide new therapeutic strategies to improve treatment response and survival in ES-SCLC.

In this study, we identify reciprocal signaling between SCLC cells and endothelial cells in which SCLC exposure induces endothelial CXCL chemokine expression, while CXCL-CXCR2 signaling promotes SCLC migration and cancer-endothelial engagement. Using quantitative Metastasis-Originated Barcode Sequencing (MOBA-seq), we show that CXCR2 promotes metastatic seeding across multiple organs and shapes subsequent clonal outgrowth states, with prominent effects in the liver. Mechanistically, RAC1-dependent cytoskeletal remodeling functions as a downstream effector of tumor-cell CXCR2 signaling. We further show that pharmacologic CXCR2 blockade suppresses SCLC migration, endothelial interactions, and liver metastatic seeding in vivo. These findings provide a mechanistic framework for cancer-endothelial crosstalk during metastatic establishment and nominate the CXCL-CXCR2-RAC1 axis as a potential vulnerability for limiting metastatic progression in SCLC.

## Results

### Single-cell profiling reveals SCLC-induced LSEC activation marked by CXCL chemokine expression

Liver sinusoidal endothelial cells (LSECs) line the hepatic sinusoids and are among the first stromal cells encountered by circulating tumor cells (CTCs) during liver metastasis^25^. ECs have dual roles during metastatic progression and can impede invasion by acting as a physical barrier or promote colonization through establishment of a permissive pre-metastatic niche. Although single-cell RNA sequencing (scRNA-seq) has revealed extensive EC heterogeneity, dynamic state transitions of LSECs in response to cancer-endothelium interactions remain poorly defined^26, 27^. To characterize LSEC state transitions during SCLC exposure, we performed scRNA-seq on BFP-labeled primary human LSECs cultured either alone or co-cultured with mCherry-labeled H82 SCLC cells at a 2:1 ratio for 24 hours. Following co-culture, CD31□BFP□mCherry□ LSECs were isolated using fluorescence-activated cell sorting (FACS) and subjected to high-depth scRNA-seq (**Figure 1a**). Mapping to the MacParland human liver reference confirmed enrichment for LSEC identities with minimal hepatocyte contamination (**Figure S1a-c**). Across one monoculture and two co-culture replicates, we profiled 41,278 cells with an average sequencing depth of 19,427 UMIs per cell. To define transcriptionally distinct LSEC subpopulations, we performed Uniform Manifold Approximation and Projection (UMAP) dimensionality reduction and unsupervised clustering, which resolved five major LSEC populations (**Figure 1b**). Comparison of cluster compositions between monoculture and co-culture conditions revealed a marked redistribution of LSEC states upon SCLC exposure: the proportions of cells in EC2 and EC4 increased, whereas EC1 and EC3 decreased. Notably, EC4 represented an SCLC-induced LSEC population that expanded from <1% in monoculture to ∼15% following co-culture (**Figure 1c**). To determine the origin of this EC4 state, we performed RNA velocity analysis to reconstruct transcriptional trajectories among LSEC clusters. Velocity vectors consistently supported a transition toward EC4 from EC1 and EC3, suggesting that SCLC exposure drives some LSECs to adopt a distinct, induced activation state (**Figure 1d**).

**Figure 1:**
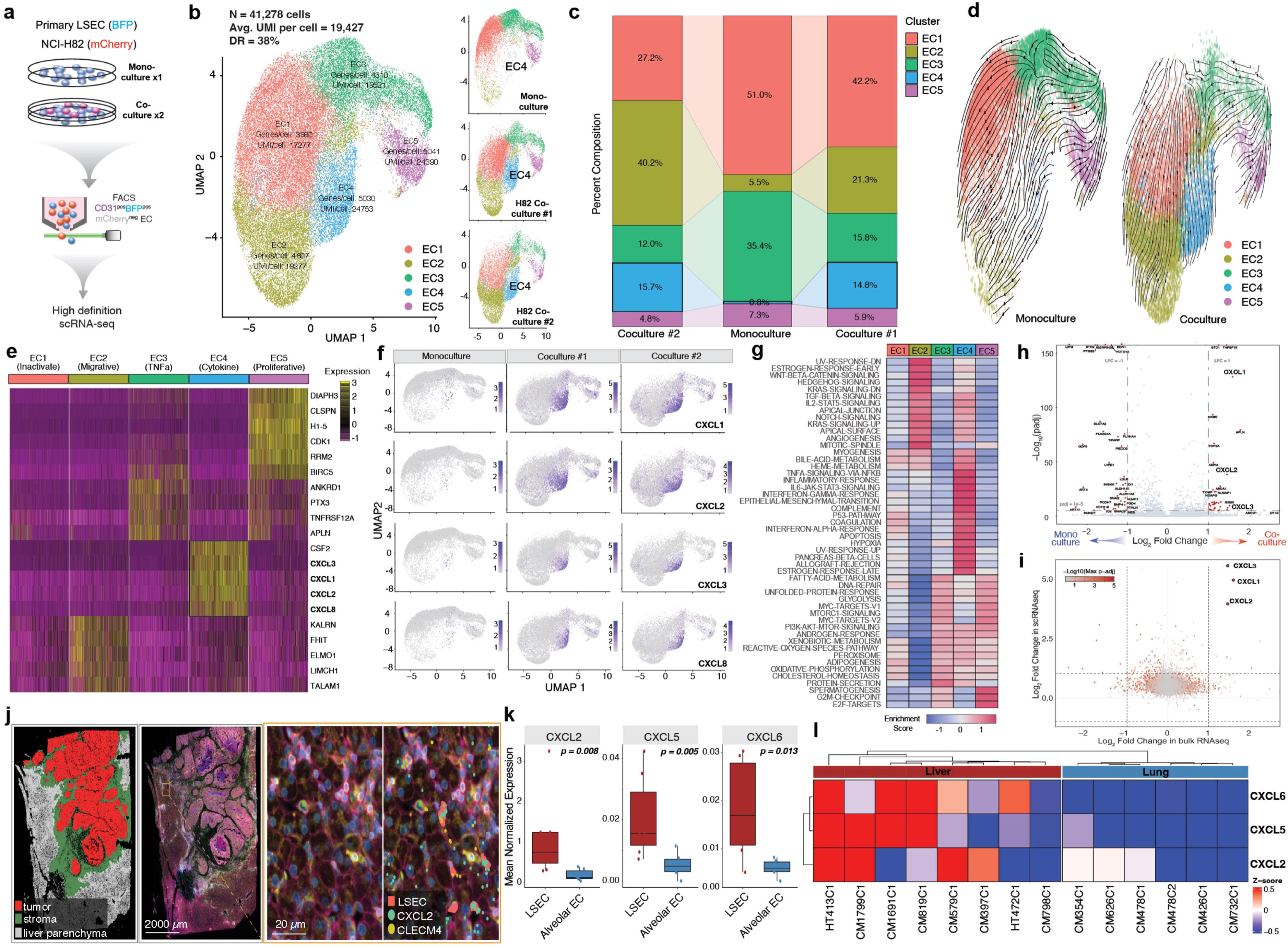
SCLC induces endothelial activation and upregulation of CXCL chemokines. **a.** Schematic illustrating isolation of monocultured and SCLC co-cultured liver sinusoidal endothelial cells (LSECs) for single-cell RNA sequencing (scRNA-seq). Primary human LSECs labeled with BFP were cultured alone or co-cultured for 24 hours with mCherry-labeled H82 SCLC cells at a 2:1 ratio. CD31+BFP+mCherry-LSECs were isolated by flow cytometry and subjected to scRNA-seq. **b.** scRNA-seq identifies distinct transcriptional states in SCLC-exposed LSECs. *Left*: UMAP visualization of five LSEC clusters (EC1-EC5) derived from monoculture and two independent H82 co-culture replicates. *Right*: split UMAPs showing redistribution of LSEC states after SCLC exposure. **c.** SCLC co-culture alters LSEC state composition. Alluvial plots show changes in the proportions of EC1-EC5 across monoculture and two co-culture replicates, including expansion of EC2 and EC4 populations. **d.** RNA velocity analysis suggests directional transitions among LSEC states after SCLC exposure. Streamlines overlaid on UMAP embeddings show trajectories toward the induced EC4 state. **e.** LSEC clusters exhibit distinct transcriptional programs. Heatmap of representative marker genes highlights quiescent, migratory, inflammatory, cytokine-high, and proliferative endothelial states. **f.** SCLC co-cultured LSECs show strong CXCL chemokine expression. Feature plots map CXCL1, CXCL2, CXCL3, and CXCL8 expression across monoculture and co-culture UMAPs. **g.** Pathway analysis identifies distinct functional programs across LSEC states. Heatmap shows normalized enrichment scores for representative gene sets, with EC4 enriched for inflammatory and stress-response programs. **h.** CXCL1 and CXCL2 are strongly upregulated in HUVECs co-cultured with H82 cells. Volcano plot shows differentially expressed genes in HUVEC monoculture versus HUVEC-H82 co-culture. **i.** CXCL1, CXCL2, and CXCL3 are concordantly induced across LSEC and HUVEC models. Scatter plot compares transcriptomic fold changes in LSECs and HUVECs after H82 SCLC co-culture. **j.** Representative spatial transcriptomic map from a colorectal cancer liver metastasis showing tumor, stroma, liver parenchyma, LSECs, CXCL2, and CLEC4M localization, including higher-resolution views of the tumor-liver interface. **k.** CXCL2, CXCL5, and CXCL6 expression is enriched in liver LSECs relative to lung alveolar ECs across the metastatic cohort. Box plots show normalized expression for each chemokine. **l.** Heatmap showing CXCL2, CXCL5, and CXCL6 expression across eight CRC liver metastasis and six CRC lung metastasis samples, demonstrating higher expression in liver ECs than lung ECs in this cohort.

To characterize transcriptional programs defining LSEC subpopulations, we performed differential gene expression (DGE) analysis across clusters. EC1 lacked distinguishing marker genes, consistent with a relatively quiescent phenotype, and was designated as inactive ECs. EC2 showed enriched expression of genes associated with cell migration, including ELMO1, LIMCH1, and TALAM1. EC3 was characterized by elevated expression of TNF signaling-related genes, such as PTX3, ANKRD1, and TNFRSF12A. Cluster EC5 exhibited enrichment of cell proliferation-related genes, including CLSPN, CDK1, and RRM2. The induced EC4 population, however, was distinguished by robust expression of multiple CXCL chemokines, including CXCL1, CXCL2, CXCL3, and CXCL8. Based on this transcriptional signature, we designated EC4 as cytokine-releasing LSECs (**Figure 1e**, **Figure S1d**). Violin plot analysis demonstrated a 10-to 100-fold increase in CXCL chemokine expression in EC4 compared with all other LSEC populations (**Figure S1e**). Consistent with this finding, visualization of CXCL expression across monoculture and co-culture conditions revealed that induction of CXCL1, CXCL2, CXCL3, and CXCL8 was restricted to the EC4 cluster and occurred exclusively following SCLC co-culture (**Figure 1f**). To define the functional programs underlying each LSEC state, we performed Gene Set Enrichment Analysis (GSEA). Inflammatory signaling, hypoxia, and epithelial-to-mesenchymal transition pathways were strongly enriched in EC4, whereas cell cycle-associated pathways were comparatively suppressed (**Figure 1g**). These results collectively indicate that SCLC-LSEC interactions drive a distinct EC state transition characterized by the emergence of a CXCL-high, cytokine-releasing LSEC state.

### SCLC induces CXCL chemokine expression in both human and mouse ECs

To determine whether SCLC-induced CXCL expression represents a general feature of SCLC-EC interactions, we performed bulk RNA sequencing of human umbilical vein endothelial cells (HUVECs) cultured alone or co-cultured with H82 SCLC cells. DGE analysis identified 742 genes that were significantly upregulated in the co-culture condition relative to monoculture, with CXCL1 and CXCL2 among the most highly induced transcripts (**Figure 1h**, **Figures S1f-i**). We next cross-compared differentially expressed genes from the HUVEC bulk RNA-seq DGE dataset with the LSEC scRNA-seq DGE dataset. Notably, CXCL1, CXCL2, and CXCL3 were consistently upregulated in both endothelial systems, indicating a shared SCLC-induced transcriptional response driving CXCL expression across distinct EC types (**Figure 1i**).

To corroborate these findings in a murine system, we cultured primary mouse LSECs alone or with mouse-derived KP-1 SCLC cells (**Figure S2a-b**). After 24 hours of co-culture, Cxcl1 and Cxcl2 mRNA levels were significantly increased in mouse LSECs relative to monocultured controls, together with additional transcripts associated with endothelial activation and cell-cell interactions (**Figure S2c**). To confirm SCLC-induced CXCL upregulation at the protein level, we performed immunofluorescence staining of CXCL1 in HUVECs co-cultured with H82 SCLC cells for 48 hours. CXCL1 expression was markedly increased in HUVECs relative to monoculture controls (**Figure S2d**). Finally, to visualize endothelial CXCL responses in vivo, we stained brain, liver, and lung tissues for CXCL1 and CD31 48 hours after intravenous transplantation of RP48 SCLC cells. CXCL1 was not detectably increased in brain or lung vasculature but was significantly elevated in liver CD31+ endothelial cells (**Figure S2e**). Together, these data identify an SCLC-induced CXCL-high endothelial state and show that the response is conserved across human and mouse endothelial models, with a particularly strong liver endothelial response *in vivo*.

### CXCL chemokine expression is enriched in liver endothelium in human metastatic samples

To examine endothelial CXCL expression in human metastatic tissues, we reanalyzed an unpublished spatial transcriptomics dataset from colorectal cancer (CRC) patients with liver and lung metastases (Peng et al., unpublished data). Endothelial subtype analysis identified LSECs as the predominant endothelial population in liver parenchyma and alveolar ECs as the dominant population in lung parenchyma (**Figures S1j-k**). A representative CRC liver metastasis showed spatial co-expression of CXCL2 with the LSEC marker CLEC4M in liver parenchyma (**Figure 1j**). Across eight liver and six lung metastasis samples, CXCL2, CXCL5, and CXCL6 expression was significantly higher in liver LSECs than in lung alveolar ECs (**Figure 1k**), and heatmap analysis showed consistent enrichment of these CXCL transcripts in liver ECs across samples (**Figure 1l**). These data support enrichment of CXCL chemokine expression in liver LSECs relative to lung alveolar ECs in this metastatic cohort.

### CXCL chemokines stimulate SCLC migration

CXCL chemokines are well established in promoting immune cell recruitment during inflammation and cancer immune responses^28, 29^. However, their roles in SCLC-endothelial interactions during metastasis remain understudied. To systematically evaluate the contribution of endothelial-derived CXCL chemokines to SCLC metastatic behaviors, we first assessed whether CXCLs affected SCLC proliferation using a CCK-8 viability assay. Compared to vehicle control, cultures containing recombinant CXCL1, CXCL2, or CXCL3 (30 ng/mL) for 72 hours did not alter the viability of either human H82 or mouse RP48 SCLC cells in vitro (**Figure 2a**). We next tested whether CXCL chemokines promoted SCLC migration using transwell assays, in which recombinant CXCLs were added to the lower chamber and migration was quantified by measuring the cell number in the lower chamber. Among the six conserved CXCL chemokines across human and mouse, CXCL1, CXCL2, and CXCL3 elicited the most robust migratory responses in non-adherent H82 and RP48 cells relative to controls (**Figure 2b**). We confirmed this finding using a combined CXCL1/2/3 treatment (“CXCLs”) in adherent H446 and DMS-273 SCLC cells, which similarly increased transwell migration (**Figure 2c**).

**Figure 2:**
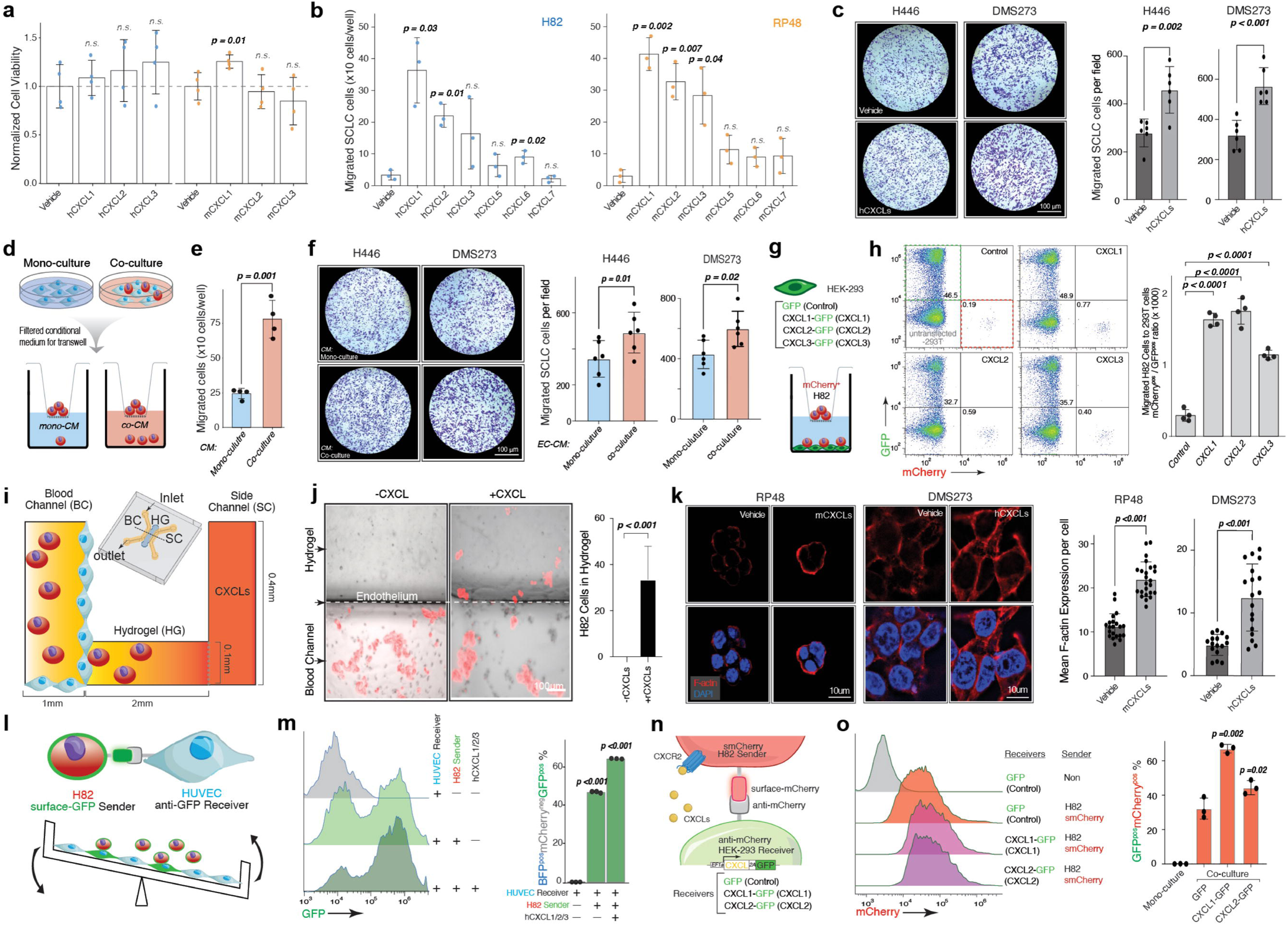
CXCL chemokines induce SCLC migration and strengthen SCLC-EC interactions. **a.** CXCL chemokines do not affect SCLC cell viability. Viability of H82 and RP48 SCLC cells treated with recombinant human or mouse CXCL1, CXCL2, or CXCL3 was quantified using a CCK-8 assay. Data are presented as mean ± SD of OD□□□fold change relative to vehicle-treated controls (n = 4 wells). **b.** CXCL chemokines enhance migration of non-adherent SCLC cells. Recombinant CXCL1, CXCL2, or CXCL3 was placed in the lower chamber, and non-adherent H82 (*Left*) or RP48 (*Right*) SCLC cells were seeded in the upper chamber. Migrated cells were quantified from 10% of the lower chamber and presented as mean ± SD (n = 3 wells). **c.** CXCL chemokines enhance migration of adherent SCLC cells. *Left:* Representative images showing migration of adherent H446 and DMS-273 SCLC cells toward vehicle control or CXCL chemokines in the lower chamber. *Right:* Migrated cells per well are quantified and reported as mean ± SD (n = 6 wells). **d.** Schematic of SCLC migration toward conditioned media. HUVEC monoculture-conditioned medium (mono-CM) or HUVEC-H82 co-culture-conditioned medium (co-CM) was placed in the lower transwell chamber, and mCherry-labeled H82 cells in the upper chamber were assessed for migration after 48 hours. **e.** HUVEC-H82 co-culture-conditioned medium promotes SCLC migration. Co-CM yielded a higher number of migrated SCLC cells relative to mono-CM. Migrated cells were quantified from 10% of the lower chamber and presented as mean ± SD (n = 4 wells). **f.** HUVEC-H82 co-culture-conditioned medium promotes transwell migration of adherent SCLC cell lines. Left: Representative images of H446 and DMS-273 migration toward mono-CM or co-CM. Right: Migrated cells per well are quantified and reported as mean ± SD (n = 6 wells). **g.** Schematic of HEK-293-SCLC co-culture transwell migration assay. GFP HEK-293 cells overexpressing CXCL1, CXCL2, or CXCL3 were placed in the lower chamber of a transwell system, and mCherry□ H82 SCLC cells were seeded in the upper chamber. Migration was quantified by flow cytometry as the percentage of mCherry□ H82 cells detected in the lower chamber relative to GFP□ HEK-293 cells. **h.** CXCL-expressing HEK-293 cells enhance SCLC migration independent of endothelial cell identity. Flow cytometry analysis (*Left*) and quantitative assessment (*Right*) show increased migration of H82 cells toward HEK-293 cells overexpressing CXCL1, CXCL2, or CXCL3, compared with control HEK-293 cells. Migration is normalized to the number of GFP HEK-293 cells in the lower chamber. Ratios of mCherry□ H82 to GFP□ HEK-293 cells per well are reported as mean ± SD (n = 4 wells). **i.** Schematic of a 3D microfluidic SCLC-EC co-culture system mimicking trans-endothelial migration. The blood channel (BC) is lined with HUVECs and contains mCherry-labeled SCLC cells, while the side channel (SC) is filled with medium supplemented with CXCL chemokines. The two channels are separated by a hydrogel (HG) matrix. SCLC trans-endothelial migration is assessed by quantification of SCLC cells that traverse the HUVEC-lined barrier and enter the hydrogel toward CXCLs in the side channel. The entire device is maintained under orbital shaking to generate physiologically relevant shear stress. **j.** CXCL chemokines stimulate H82 trans-endothelial migration. Left: Representative images showing mCherry-labeled H82 cells migrating into the hydrogel in the absence or presence of CXCL chemokines. Right: Quantification of H82 trans-endothelial migration. CXCL supplementation increased SCLC trans-endothelial migration into the hydrogel. Data reported as mean ± SD of mCherry+ H82 per chip (n = 3 chips). **k.** CXCLs induce F-actin assembly in SCLC cells. *Left*: Representative immunofluorescence images of F-actin staining in RP48 and DMS-273 SCLC cell lines treated with or without CXCL chemokines. *Right*: Quantification of F-actin signal per cell, presented as mean ± SD of the F-actin to DAPI signal intensity ratio per field (n > 15 fields). CXCL treatment significantly increased F-actin assembly in both SCLC cell lines. **l.** Schematic of quantification of SCLC-EC interactions via the G-baToN system. H82 SCLC cells (mCherry□) expressing cell surface GFP served as sender cells, while HUVECs (BFP) expressing cell surface anti-GFP served as receiver cells. GFP transfer during direct cell-cell contact was assessed by flow cytometry, and cultures were subjected to orbital shaking to model physiological shear stress. **m.** CXCL chemokines strengthen interactions between H82 SCLC cells and HUVECs. *Left:* Flow cytometry analysis of GFP transfer in monocultured HUVEC receivers and in co-cultures of H82 senders with HUVEC receivers, with or without CXCLs supplementation. *Right*: Quantification of SCLC-EC interaction strength, measured as the percentage of GFP cells within the viable BFP mCherry□ HUVEC receiver population (mean ± SD; N = 3). CXCL treatment significantly increased GFP transfer into HUVECs, indicating enhanced physical contact between H82 cells and HUVECs. **n.** Schematic of quantification of CXCL-dependent effects on SCLC-HEK-293 interactions. Cell surface mCherry-expressing H82 SCLC cells (senders) and anti-mCherry-expressing HEK-293 cells (receivers) were co-cultured with or without additional overexpression of CXCL chemokines, under orbital shaking to mimic physiological shear stress. Direct H82-HEK-293 contact enabled mCherry□ transfer to HEK-293 receiver cells, which was quantified by flow cytometry as a measure of interaction strength. **o.** CXCL overexpression strengthens interactions between H82 cells and HEK-293 cells. *Left*: Flow cytometry analysis of mCherry□ transfer in monocultured HEK-293 receiver cells and in co-cultures of H82 sender cells with HEK-293 receivers expressing GFP control, CXCL1, or CXCL2. *Right*: Quantification of H82-HEK-293 interaction strength, measured as the percentage of mCherry cells within the viable GFP□ HEK-293 receiver population (mean ± SD; N = 3). CXCL overexpression significantly increased HEK-293 interaction with SCLC independently of endothelial identity.

Having established that SCLC exposure induces endothelial CXCL expression and that recombinant CXCL chemokines promote SCLC migration, we next asked whether soluble factors present in SCLC-HUVEC co-culture-conditioned medium were sufficient to enhance tumor-cell migration. We collected conditioned medium from HUVEC monocultures (mono-CM) or HUVEC-H82 co-cultures (co-CM) and used these media as chemoattractants in the lower chamber of transwell assays (**Figure 2d**). H82 cells migrated more efficiently toward co-CM than toward mono-CM, and the same effect was observed in adherent H446 and DMS-273 cells (**Figure 2e-f**). These experiments show that soluble factors present in SCLC-HUVEC co-culture-conditioned medium enhance SCLC migration, although they do not by themselves assign the source of those factors to a specific cell type. We next tested whether CXCL expression alone was sufficient to increase SCLC migration independent of endothelial identity by engineering HEK-293 cells to express GFP control or human CXCL1, CXCL2, or CXCL3 and placing them in the lower chamber of a transwell system. Migration of mCherry-labeled H82 cells toward the HEK-293 cells was quantified by flow cytometry after 24 hours (**Figure 2g**). CXCL-expressing HEK-293 cells significantly increased H82 migration relative to control HEK-293 cells (**Figure 2h**), demonstrating that CXCL signaling is sufficient to enhance SCLC migration in this assay.

In the metastatic cascade, cancer cells must directly interact with endothelial cells during trans-endothelial migration (TEM), a key step in both intravasation and extravasation. With the aim of assessing the role of CXCL chemokines in TEM under more physiological conditions, we employed a 3D microfluidic co-culture system to mimic the perivascular microenvironment^30^. The platform consisted of a blood channel lined with HUVECs and containing freely flowing mCherry-labeled H82 cells, separated from an adjacent side channel by a hydrogel matrix. Shear stress was applied by a rocking shaker to induce SCLC movement in the blood channel and increase HUVEC interaction. (**Figure 2i**). Addition of CXCL chemokines to the side channel significantly increased TEM of H82 cells across the HUVEC barrier and into the hydrogel compared with unstimulated controls (**Figure 2j**). Given that both CXCL chemokines and the actin cytoskeleton are key drivers of cell motility, we next asked whether CXCL stimulation promotes F-actin assembly in SCLC cells. Phalloidin staining revealed that CXCL treatment significantly increased F-actin polymerization in both RP48 and DMS-273 cells (**Figure 2k**), consistent with CXCL-induced activation of a pro-migratory cytoskeletal program.

### CXCL chemokines strengthen SCLC-EC interactions

SCLC trans-endothelial migration requires direct cancer-endothelial interactions at the vascular wall during metastasis. To directly test whether CXCLs enhance cancer-endothelial adhesion, we used the G-baToN system to quantitatively measure physical cell-cell contact^30^. In this assay, “sender” SCLC cells express a membrane-tethered fluorescent protein (e.g., GFP or mCherry), and “receiver” endothelial cells express a surface nanobody against that fluorescent protein. Direct contact between sender and receiver cells enables fluorescent protein transfer into receiver cells in a time-dependent quantitative manner allowing interaction strength to be measured by flow cytometry. We generated mCherry-labeled H82 cells as GFP senders and BFP-labeled HUVECs as anti-GFP receivers. We co-cultured senders and receivers at a 1:1 ratio with or without CXCL chemokines under orbital shaking (**Figure 2l**). After 24 hours, addition of CXCLs significantly increased the fraction of GFP□ HUVEC receiver cells, indicating enhanced physical interactions between SCLC cells and endothelial cells (**Figure 2m**). To determine whether this CXCL-dependent cell-cell interaction is specific to endothelial cells, we performed an analogous G-baToN assay using H82 mCherry sender cells and HEK-293 anti-mCherry receiver cells engineered to overexpress either control or CXCL1 or CXCL2 (**Figure 2n**). HEK-293 cells overexpressing CXCL1 or CXCL2 exhibited increased mCherry uptake compared with control HEK-293 cells, supporting CXCL-dependent enhancement of SCLC-stromal cell interactions independent of endothelial identity (**Figure 2o**). Together, these results indicate that CXCL chemokines strengthen SCLC-stromal cell interactions, consistent with a model in which CXCL signaling can strengthen SCLC-stromal interactions and may thereby facilitate tumor-cell engagement with the vascular wall.

### CXCR2 signaling promotes metastatic seeding and shapes metastatic outgrowth states

CXCR2 is the shared receptor for the CXCL1/2/3 chemokines, and its expression has been associated with prognostic significance across multiple cancer types^31, 32^. However, therapeutic targeting of CXCR2 clinically has yielded unsatisfactory outcomes, likely reflecting an incomplete understanding of its context-dependent roles and therapeutic window during cancer progression and metastasis. Metastasis is a multistep process that, within target organs, encompasses extravasation, metastatic seeding, transition to metastatic outgrowth, and clonal expansion^9, 10^. To precisely delineate the function of CXCR2 across distinct stages of the SCLC metastatic cascade, we developed Metastasis Originated BArcode Sequencing (MOBA-seq), a quantitative approach assessing gene function throughout the metastatic cascade at single-colony resolution^33^. We transduced Cas9-expressing RP48 SCLC cells with a randomly barcoded lentiviral sgRNA library containing a non-targeting control sgRNA (*sgCtrl*) and three independent sgRNAs targeting *Cxcr2* (*sgCxcr2*). Barcoded SCLC cells were then intravenously transplanted into *NOD/Scid/IL2R*_γ_*-null (NSG)* mice, and metastasis-bearing tissues were harvested at 2 days, 1 week, 2 weeks, and 3 weeks post-injection for barcode sequencing and MOBA-seq analysis (**Figure 3a**).

**Figure 3:**
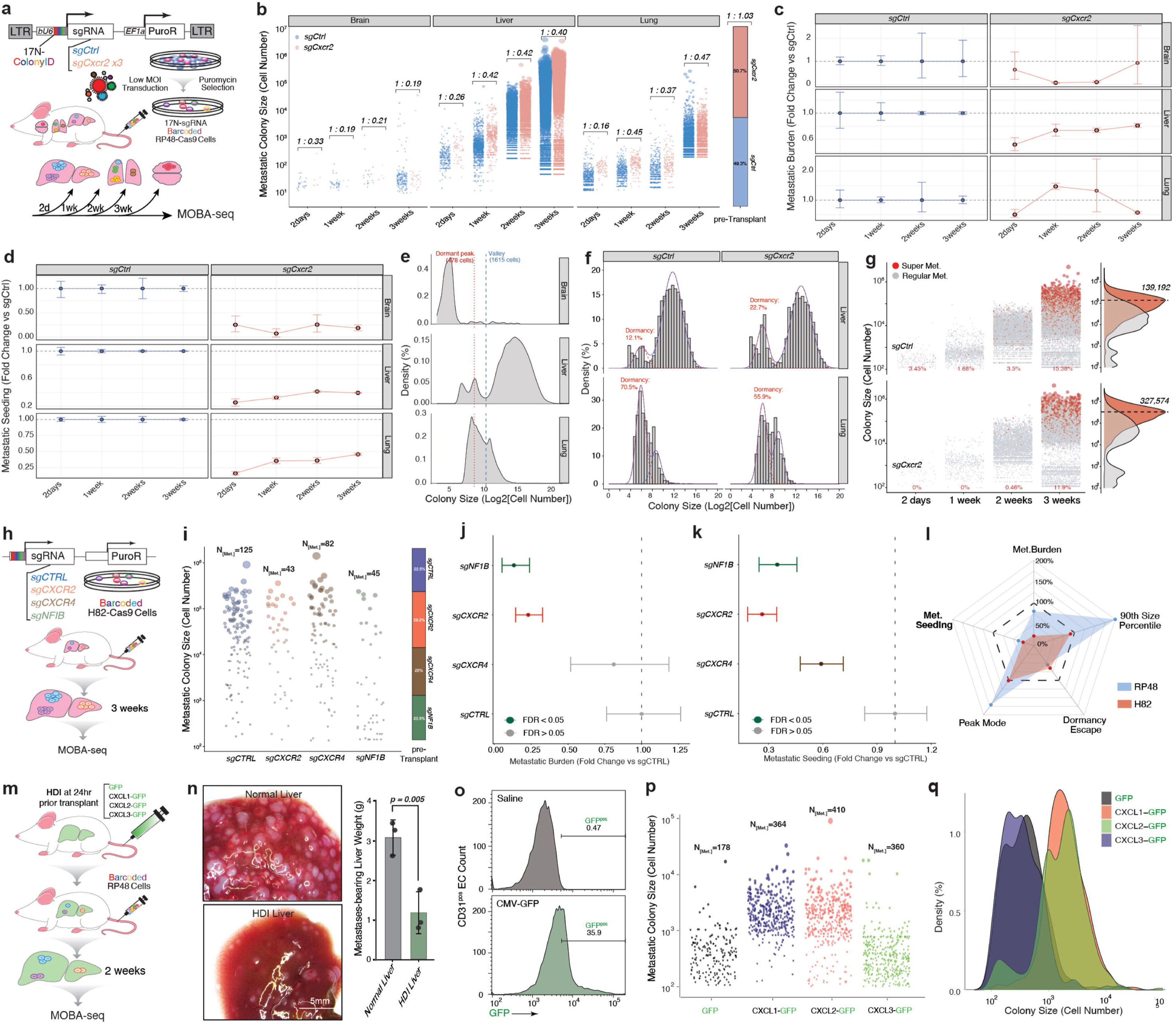
Quantitative dissection of CXCR2 signaling in SCLC liver metastases. **a.** Schematic of quantitative assessment of CXCR2 function in SCLC metastasis using Metastasis-Originated Barcode Sequencing (MOBA-seq). Cas9-expressing RP48 SCLC cells were randomly barcoded with a lentiviral sgRNA library containing one control sgRNA and three sgRNAs targeting *Cxcr2*. Barcoded SCLC cells were subsequently transplanted intravenously into recipient mice, and metastatic tissues were harvested at multiple time points. Genomic DNA from metastatic lesions in target organs was subjected to barcode sequencing and analyzed using the MOBA-seq pipeline to quantify genotype-specific contributions to metastatic seeding and outgrowth. **b.** *Cxcr2* inactivation suppresses SCLC metastasis across multiple tissues and time points. Metastatic colonies recovered from mouse brain, liver, and lung at four time points were plotted, with each dot representing an individual barcode-derived metastatic colony and dot size scaled to colony cell number. Across all organs and time points, *Cxcr2* inactivation reduced the number of metastatic colonies compared with controls. Pre-transplantation barcode distributions showed similar cell numbers between the two genotypes. **c.** *Cxcr2* inactivation suppresses SCLC metastatic burden across multiple tissues and time points. Metastatic burden in brain, liver, and lung was quantified via MOBA-seq at 2 days, 1 week, 2 weeks, and 3 weeks post-transplantation. For each tissue and time point, relative metastatic burden was plotted as fold change compared with the control sgRNA. Data points outlined in black denote comparisons with statistical significance (FDR < 0.05). **d.** *Cxcr2* inactivation suppresses SCLC metastatic seeding across multiple tissues and time points. Relative metastatic seeding was quantified in brain, liver, and lung at 2 days, 1 week, 2 weeks, and 3 weeks post-transplantation. Fold changes in metastatic seeding were plotted relative to sg*Ctrl*. Data points outlined in black denote statistical significance (FDR < 0.05). Cxcr2 inactivation led to a marked reduction in the number of seeded metastatic colonies across all tissues and time points. **e.** MOBA-seq resolves small, non-expanding and expanding metastatic colony states across tissues. The distribution of log -transformed metastatic colony sizes from liver, lung, and brain at the 3-week time point was modeled to identify two key features: a small-colony mode (left mode, red) and a valley mode (right mode, blue) demarcating the transition from small, non-expanding to expanding colonies. These empirically derived cutoff values were uniformly applied across all three tissues to enable unbiased quantification of genotype-specific effects on dormant-like metastatic states. Notably, RP48 cells showed a liver-enriched shift from the small-colony mode toward larger expanding colonies. **f.** *Cxcr2* inactivation increases the dormant-like, non-expanding colony fraction in the liver, but not the lung, at 3 weeks. A Gaussian mixture model was applied to the log -transformed metastatic colony size distributions from liver and lung, using the dormant-mode and valley-mode cutoffs defined in (e) to quantify genotype-specific occupancy of the dormant-like, non-expanding state in each organ. In the liver, Cxcr2 inactivation substantially increased the proportion of dormant-like colonies compared with control. In contrast, Cxcr2 inactivation led to a reduction in dormant-like colonies in the lung. **g.** *Cxcr2* inactivation suppresses dissemination of metastatic SCLC cells into the blood. Shared barcodes detected in both liver metastases and circulating tumor cells were used to identify metastatic colonies capable of further dissemination (“SuperMets,” highlighted in red). *Left*: Jitter plots of colony sizes across time points show the frequency and distribution of SuperMet colonies for each genotype. *Right*: Size distributions of SuperMet versus regular metastatic colonies. *Cxcr2* inactivation reduced the number of SuperMet colonies and increased the colony size threshold for dissemination. **h.** Schematic showing MOBA-seq comparison of the metastatic effects of *CXCR2* and established metastatic drivers in human SCLC. Barcoded H82-Cas9 SCLC cells were transduced with a lentiviral sgRNA library containing a control sgRNA and sgRNAs targeting *CXCR2*, *CXCR4*, or *NFIB*. After intravenous transplantation of the barcoded pools into recipient mice, metastasis-bearing liver tissues were harvested for library preparation, sequencing, and analysis. MOBA-seq was performed to quantify genotype-specific metastatic fitness and evaluate the relative effect of *CXCR2* compared with known metastatic drivers *CXCR4* and *NFIB*. **i.** *CXCR2* inactivation suppresses liver metastasis at 3 weeks, comparable to inactivation of other metastatic driver genes. Metastatic colonies recovered from liver at 3 weeks were plotted, with each dot representing a single barcode-derived colony and dot size scaled to colony cell number. Inactivation of *CXCR2*, *CXCR4*, or *NFIB* each resulted in a reduction in metastatic colony number relative to control. Pre-transplantation barcode distributions confirmed similar input cell numbers across all genotypes. **j.** *CXCR2* inactivation suppresses metastatic burden as strongly as *NFIB*. Relative metastatic burden in the liver at 3 weeks was plotted as fold change compared with control sgRNA, with statistically significant differences (FDR < 0.05) shown in color. Both *CXCR2* and *NFIB* inactivation resulted in significantly reduced metastatic burden. **k.** *CXCR2* inactivation suppresses metastatic seeding more strongly than other metastatic driver genes. Relative metastatic seeding in the liver at 3 weeks was plotted as fold change compared with control sgRNA, with statistically significant differences (FDR < 0.05) shown in color. *CXCR2* inactivation produced the greatest reduction in metastatic seeding, exceeding the effects observed with *CXCR4* or *NFIB* inactivation. **l.** *CXCR2*-dependent metastatic phenotypes are conserved between human and mouse SCLC cell lines. Radar plots summarizing liver metastatic metrics for *Cxcr2* demonstrate similar suppression of metastatic burden, metastatic seeding, and dormancy escape in human H82 and mouse RP48 SCLC cell lines. Notably, inactivation of *Cxcr2* in RP48 cells results in higher 90th-percentile colony sizes and peak mode values compared with H82 cells, indicating cell line-specific CXCR2 effects on clonal expansion dynamics. **m.** Schematic of quantification of CXCL chemokines as pro-metastatic factors in SCLC liver metastasis via MOBA-seq. Mice were treated by hydrodynamic injection (HDI) to deliver plasmids expressing either GFP alone (control) or GFP-tagged CXCL chemokines. 24 hours after HDI, barcoded SCLC cells were transplanted intravenously and metastasis-bearing livers were harvested two weeks later. MOBA-seq was then performed to quantify the impact of CXCL overexpression on liver metastatic outgrowth. **n.** Hydrodynamic injection suppresses baseline SCLC liver metastasis. *Left*: Representative images of metastasis-bearing mouse livers with and without HDI treatment show a marked reduction in H82 SCLC metastatic lesions in HDI-treated livers. *Right*: Quantification of metastasis-bearing liver weight per mouse demonstrates a significant decrease in metastatic burden following HDI treatment, presented as mean ± SD (N = 3 mice). **o.** Hydrodynamic injection efficiently transfects liver ECs. Flow cytometry analysis of GFP expression in liver CD31^+^ EC following HDI with either saline or CMV-GFP plasmid. HDI achieved robust transfection, with 35.9% of liver endothelial cells expressing GFP. **p.** CXCL overexpression increases SCLC metastatic seeding in HDI-treated mouse livers. Metastatic colonies recovered from HDI-treated mouse livers were plotted, with each dot representing an individual barcode-derived colony and dot size scaled to colony cell number. Overexpression of CXCL1, CXCL2, or CXCL3 increased the number of metastatic colonies relative to GFP control. **q.** CXCL1 and CXCL2 overexpression shift SCLC metastases toward larger, expanding colonies. The log -transformed metastatic colony size distributions for GFP control and CXCL-overexpressing conditions were plotted. CXCL1- and CXCL2-overexpressing livers showed a shift toward expanding metastatic colonies compared with the distribution enriched for small, non-expanding colonies observed in GFP control livers.

Across tissues and time points, Cxcr2-targeted cells produced fewer metastatic colonies than controls despite nearly identical representation in the pre-transplantation cell pools, consistent with impaired extravasation and/or seeding (**Figure 3b**). Cxcr2 inactivation also reduced total metastatic SCLC cell number across multiple tissues and time points (**Figure 3c**). Because metastatic seeding is a major determinant of metastatic burden, we next quantified seeding efficiency and found that Cxcr2 inactivation reduced metastatic seeding in all tissues examined at every time point analyzed (**Figure 3d**). We then examined subsequent clonal behavior using colony-size distributions (**Figures S3a-b**). Pooling colonies across tissues revealed a small-colony mode consistent with a dormant-like, non-expanding state and a valley separating this mode from expanding colonies (**Figure 3e**). Using empirically derived cutoffs and a Gaussian mixture model, we quantified the fraction of colonies occupying this dormant-like state. In the liver, Cxcr2 inactivation increased the dormant-like fraction, whereas the lung showed the opposite trend (**Figure 3f**). Finally, Cxcr2 inactivation reduced the frequency of liver SuperMet colonies capable of further dissemination into the bloodstream and increased the colony-size threshold associated with dissemination (**Figure 3g**). Together, these results support a broad role for Cxcr2 in metastatic seeding and a liver-enriched influence on the transition from small, non-expanding colonies to metastatic outgrowth.

### CXCR2-dependent metastatic seeding phenotypes are conserved across human and mouse SCLC

To validate CXCR2 function across species and benchmark its metastatic effects against established SCLC metastatic drivers, we performed MOBA-seq using barcoded human H82 SCLC cells transduced with a library containing sgCTRL, sgCXCR2, and sgRNAs targeting the known metastatic drivers CXCR4 and NFIB (**Figure 3h**)^34–36^. At three weeks post-transplantation, CXCR2 inactivation reduced liver metastatic seeding to a degree comparable to NFIB inactivation (**Figure 3i**). Analysis of liver metastatic burden showed that CXCR2 and NFIB inactivation produced the strongest suppression of overall metastatic burden (**Figure 3j**), while direct comparison of seeding efficiencies showed the largest reduction with CXCR2 inactivation (**Figure 3k**). Cross-species comparison demonstrated broadly concordant effects of CXCR2 inactivation on metastatic burden and seeding, with cell-line-specific differences in colony-size and outgrowth metrics (**Figure 3l**; **Figures S3c-f**). Together, these results indicate that the requirement for CXCR2 in metastatic seeding is conserved across human and mouse SCLC models.

### Liver CXCL overexpression increases SCLC metastatic seeding and clonal outgrowth

To test whether increased hepatic CXCL expression is sufficient to influence SCLC metastatic seeding and outgrowth, we used hydrodynamic injection (HDI) to express GFP control or GFP-tagged CXCL1, CXCL2, or CXCL3 in the liver before SCLC transplantation^37, 38^. NSG mice received HDI followed 24 hours later by tail-vein transplantation of barcoded RP48 cells, and livers were collected two weeks later for MOBA-seq analysis (**Figure 3m**). HDI itself reduced baseline metastatic burden, consistent with liver injury caused by the procedure (**Figure 3n**), and GFP expression was detected in a substantial fraction of liver CD31+ endothelial cells after HDI (**Figure 3o**). Within this HDI-treated context, expression of CXCL1, CXCL2, or CXCL3 increased the number of metastatic colonies relative to GFP control (**Figure 3p**). CXCL1 and CXCL2 also shifted colony-size distributions toward larger, expanding colonies (**Figure 3q**). These data support a pro-metastatic effect of increased liver CXCL expression on SCLC seeding and clonal outgrowth.

### CXCR2 is critical for CXCL responses in SCLC cells

Our previous results showed that the CXCR2 ligands CXCL1, CXCL2, and CXCL3 enhance SCLC migration and SCLC-EC interactions. To test whether these phenotypes are CXCR2 dependent, we first assessed baseline cell fitness by measuring viability of CXCR2-knockout (*CXCR2-KO*) human H82 and mouse RP48 cells and observed no significant differences compared with CXCR2–wild-type (*CXCR2-WT*) cells (**Figure 4a**). We then evaluated CXCR2-dependent SCLC motility in transwell assays and found that loss of CXCR2 significantly reduced migration under both baseline conditions and following CXCL stimulation (**Figure 4b**). To quantify cell migration under identical conditions, we performed a color-coded competition assay in which *CXCR2-WT* (GFP) and *CXCR2-KO* (mCherry) H82 cells were differentially labeled and allowed to compete for migration toward CXCL-containing media (**Figure 4c**). Under CXCL stimulation, *CXCR2-WT* cells consistently outcompeted *CXCR2-KO* cells in transwell migration (**Figure 4d**). Conversely, CXCR2 overexpression (*CXCR2-OvE*) did not substantially alter baseline migration but increased SCLC sensitivity to CXCL-dependent transwell migration in both non-adherent H82 cells and adherent DMS-273 and H446 cells (**Figure 4e–f**). We also examined whether CXCR2 is required for CXCL-induced F-actin assembly. CXCL stimulation increased F-actin polymerization in *CXCR2-WT* cells but not in *CXCR2-KO* cells (**Figure 4g**). Consistent with CXCR2 dependence, CXCR2 overexpression further enhanced CXCL-induced F-actin assembly (**Figure 4h**). Finally, applying the G-baToN assay, we found that CXCR2-knockout significantly decreased SCLC-EC interaction strength (**Figure 4i**). Together, these results indicate that cancer-intrinsic CXCR2 is critical for CXCL-driven migration and SCLC-EC interactions.

**Figure 4:**
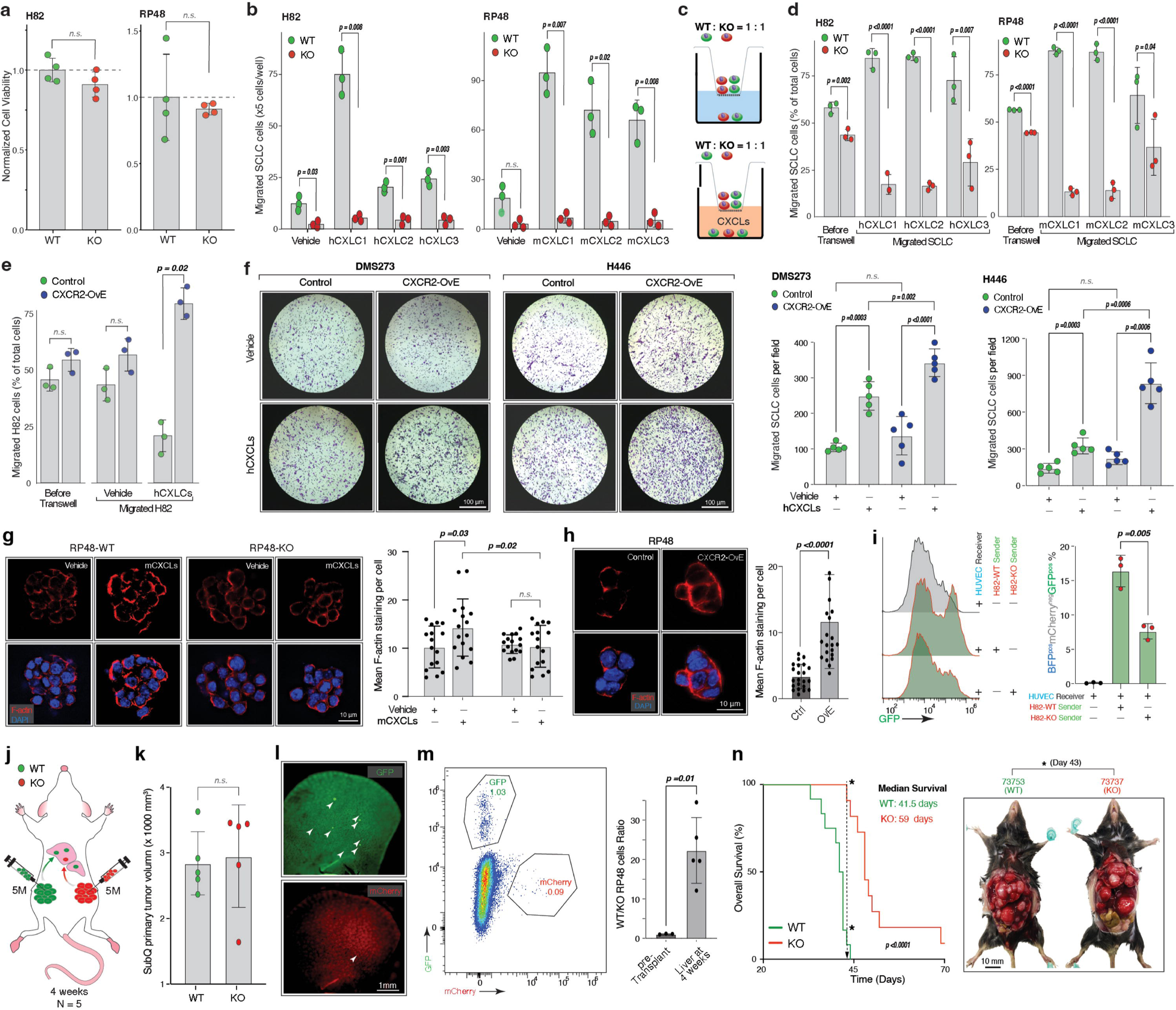
CXCR2 is required for SCLC migration and metastasis. **a.** Loss of *CXCR2* does not affect SCLC cell viability. Viability of wild-type (WT) and *CXCR2*-knockout (KO) H82 and RP48 SCLC cells was quantified using CCK-8 assay. After 72 hours of culture, no significant (n.s.) differences in cell viability were detected between WT and KO cells in either line. Data are presented as mean ± SD of OD fold change normalized to WT controls (n = 4 wells). **b.** Loss of *CXCR2* suppresses CXCL-induced SCLC cell migration. Transwell migration assays were performed by adding vehicle or CXCL1, CXCL2, or CXCL3 to the lower chamber and seeding WT or KO H82 or RP48 SCLC cells in the upper chamber. Migrated SCLC cells were quantified from 20% of the lower chamber and are presented as mean ± SD (n = 3 wells). Loss of *CXCR2* significantly reduced migration under both baseline (vehicle) conditions and all CXCL-stimulated conditions. **c.** Schematic of color-coded SCLC transwell competition assay. Equal numbers of GFP WT and mCherry□ *CXCR2*-KO SCLC cells were seeded into the upper chamber of a transwell system, with CXCL chemokines added to the lower chamber. Following migration, the proportion of GFP□ WT and mCherry□ KO cells in the lower chamber was quantified by flow cytometry to assess competitive migratory advantage between the two populations. **d.** Loss of CXCR2 impairs competitive SCLC transwell migration. GFP-labeled WT and mCherry-labeled CXCR2-KO H82 or RP48 SCLC cells were mixed at a 1:1 ratio and subjected to the transwell competition assay. Data are presented as mean ± SD of the relative percentage of GFP and mCherry cells normalized to total cells (n = 3 wells). WT cells consistently outcompeted KO cells in migration toward CXCL-containing lower chambers. **e.** CXCR2 overexpression (OvE) enhances SCLC response to CXCL-induced migration. Transwell migration of GFP-labeled control and mCherry-labeled CXCR2-OvE H82 SCLC cells was quantified by flow cytometry under vehicle or CXCL-stimulated conditions. Data are presented as mean ± SD of the relative percentage of GFP and mCherry cells normalized to total cells (n = 3 wells). CXCR2 overexpression did not alter baseline (vehicle) migration but significantly increased CXCL-driven migration. **f.** CXCR2 overexpression enhances adherent SCLC response to CXCL-induced migration. *Left*: Representative transwell images of control and CXCR2-OvE adherent SCLC cell lines (DMS-273 and H446) under vehicle or CXCL-stimulated conditions. *Right*: Quantification of migrated SCLC cells per field for each condition, presented as mean ± SD (N = 5 fields). CXCR2 overexpression did not alter baseline migration but significantly increased migration in response to CXCL treatment. **g.** *CXCR2* inactivation suppresses CXCL-induced F-actin assembly in SCLC cells. *Left*: Representative immunofluorescence images of WT and CXCR2-KO RP48 SCLC cells treated with vehicle or CXCLs, showing F-actin (red) and DAPI-stained nuclei (blue). *Right*: Quantification of F-actin levels per cell, presented as mean ± SD of the F-actin/DAPI signal intensity ratio per field (n > 15 fields). CXCL stimulation increased F-actin assembly in WT cells but not in CXCR2-KO cells. **h.** CXCR2 overexpression increases F-actin assembly in SCLC cells treated with CXCLs. *Left*: Representative immunofluorescence images of control and CXCR2-OvE RP48 SCLC cells showing F-actin (red) and DAPI-stained nuclei (blue). *Right*: Quantification of F-actin levels per cell, presented as mean ± SD of the F-actin/DAPI signal intensity ratio per field (n > 15 fields). CXCR2-OvE cells exhibited significantly enhanced F-actin organization compared with control cells. **i.** *CXCR2* inactivation in SCLC cells decreases SCLC-endothelial cell interaction strength. *Left*: Flow cytometry analysis of GFP transfer in monocultured HUVEC receiver cells and in co-cultures with WT or CXCR2-KO H82 SCLC sender cells. *Right*: Quantification of the percentage of GFP BFP mCherry^-^ HUVEC receiver cells when co-cultured with WT or CXCR2-KO H82 senders. Loss of CXCR2 significantly decreased GFP transfer into HUVEC receiver cells. **j.** Schematic of liver dissemination from paired subcutaneous SCLC tumors. Equal numbers of GFP-labeled CXCR2-WT cells and mCherry-labeled CXCR2-KO cells were injected into opposite flanks of mice to generate established subcutaneous tumors. After four weeks, both subcutaneous primary tumors and metastasis-bearing liver tissues were collected, and the relative metastatic contribution of each genotype was quantified by comparing the abundance of GFP (WT) and mCherry (KO) cells. **k.** *CXCR2* inactivation in SCLC cells does not impair primary subcutaneous tumor growth. Subcutaneous tumor volumes generated by WT and CXCR2-KO RP48 SCLC cells were quantified and are presented as mean ± SD (n = 5 mice). Primary tumor growth was comparable between WT and CXCR2-KO tumors. **l.** *CXCR2* inactivation suppresses SCLC liver metastatic colonization. Representative fluorescence images of liver surface metastases (white arrows) derived from disseminated GFP WT and mCherry□ *CXCR2*-KO RP48 SCLC cells. *CXCR2*-KO cells exhibited markedly reduced metastatic colonization of the liver. **m.** *CXCR2* inactivation reduces metastatic SCLC cell dissemination in the liver. *Left*: Flow cytometry analysis of GFP□ WT and mCherry□ CXCR2-KO RP48 SCLC cells isolated from metastasis-bearing livers four weeks after subcutaneous transplantation. *Right*: Quantification of the GFP to mCherry□ RP48 cell ratio per mouse liver, presented as mean ± SD (n = 5 mice). WT RP48 cells represented a significantly larger proportion of liver-disseminated SCLC cells compared with CXCR2-KO cells. **n.** *CXCR2* inactivation prolongs survival and reduces liver metastatic colonization. *Left*: Kaplan-Meier survival curves of mice injected via tail vein with WT or CXCR2-KO RP48 SCLC cells. Loss of CXCR2 significantly extended overall survival relative to WT controls. *Right*: Representative necropsy images from mice that succumbed on day 43 illustrate the markedly reduced liver metastatic colonization in CXCR2-KO-recipient mice.

### CXCR2 is critical for SCLC liver metastasis

To test the role of CXCR2 during dissemination from established tumors, we subcutaneously transplanted GFP-labeled CXCR2-WT and mCherry-labeled CXCR2-KO RP48 cells into opposite flanks of the same mice and allowed tumors to develop for four weeks (**Figure 4j**). CXCR2 loss did not impair primary subcutaneous tumor growth (**Figure 4k**), but representative liver imaging showed reduced metastatic colonization by CXCR2-KO cells (**Figure 4l**). Flow cytometry provided quantitative confirmation that CXCR2-WT cells comprised a significantly larger fraction of liver-disseminated RP48 cells than CXCR2-KO cells (**Figure 4m**). To determine whether CXCR2 loss also affected outcome in an experimental metastasis model, we intravenously transplanted equal numbers of CXCR2-WT or CXCR2-KO RP48 cells into C57BL/6 mice. Mice receiving CXCR2-KO SCLC cells exhibited significantly prolonged overall survival and fewer liver metastatic colonies at endpoint (**Figure 4n**). Together, these data support a requirement for CXCR2 in efficient SCLC metastatic establishment *in vivo*.

### CXCL-CXCR2 signaling promotes RAC1-dependent cytoskeletal remodeling

Our previous results established that CXCL-CXCR2 signaling promotes SCLC migration and F-actin remodeling. To identify candidate downstream signaling programs, we performed bulk RNA-seq in CXCR2-WT, CXCR2-KO, and CXCR2-overexpressing H82 cells with or without CXCL1/2/3 stimulation. Pathway-level analyses showed suppression of cytoskeletal, adhesion, and small-GTPase programs after CXCR2 loss, including Rac-, Rho-, and Rap-associated pathways (**Figure 5a**; **Figure S4a-i**). To compare the roles of Rac-, Rho-, and Rap-associated pathways in SCLC metastasis, we reanalyzed our previously reported MOBA500 screen^33^. Rac-pathway perturbations, particularly Rac1 inactivation, showed the most consistent suppression of RP48 metastatic seeding across C57BL/6 and NSG host backgrounds (**Figure 5b**; **Figure S4j**).

**Figure 5.**
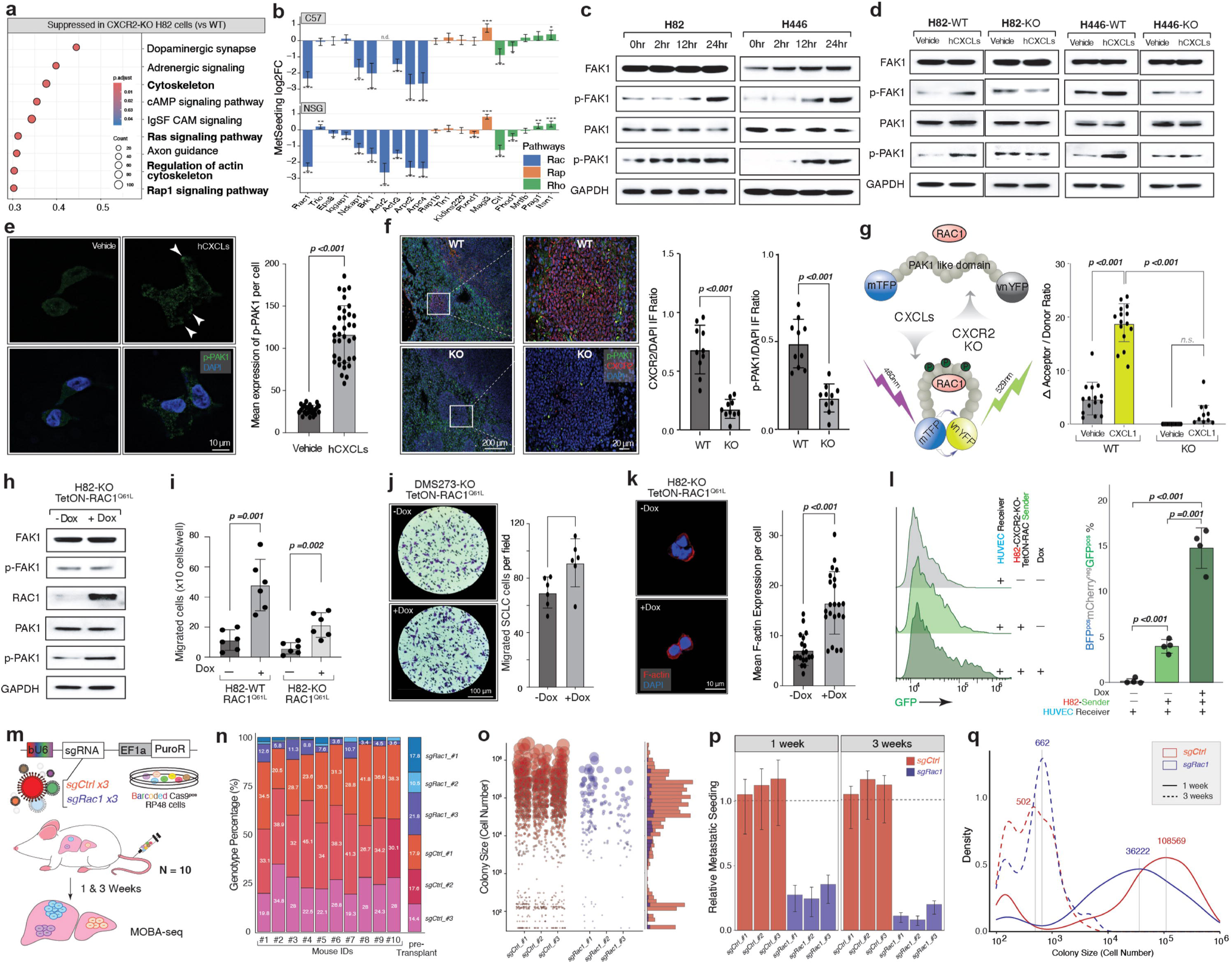
RAC1 functions downstream of CXCL-CXCR2 signaling to promote SCLC metastatic phenotypes. **a.** Transcriptomic pathway analysis identifies cytoskeletal and small-GTPase programs suppressed by CXCR2 loss. Gene-set enrichment highlights reduced cytoskeleton, Ras signaling, axon guidance, regulation of actin cytoskeleton, and Rap1 signaling pathways in CXCR2-KO relative to WT H82 cells. **b.** Re-analysis of the previously reported MOBA500 dataset^33^ showing the effects of sgRNA-mediated inactivation of genes in the Rac-, Rap-, and Rho-family signaling pathways on metastatic seeding in C57BL/6 (B6) and NSG mice. Bars indicate the log2 fold change in metastatic seeding relative to non-targeting control sgRNAs, with genes grouped by pathway. *Rac1* inactivation consistently reduced metastatic seeding in both host backgrounds. **c.** CXCL stimulation increases FAK1 and PAK1 phosphorylation in SCLC cells. Representative immunoblots of H82 and H446 cells collected at 0, 2, 12, and 24 hours after CXCL treatment show time-dependent changes in FAK1/PAK1 signaling; corresponding quantification is shown in Supplementary Figure 5a. **d.** CXCR2 loss attenuates CXCL-induced FAK1/PAK1 signaling. Representative immunoblots of CXCR2-WT and CXCR2-KO H82 and H446 cells treated with vehicle or CXCLs; corresponding quantification is shown in Supplementary Figure 5b. **e.** CXCL stimulation enriches phosphorylated PAK1 at the leading edge of adherent SCLC cells. Representative images of DMS-273 cells treated with vehicle or CXCLs show p-PAK1 (green) and DAPI (blue); arrows indicate leading-edge signal. Right: quantification of p-PAK1 signal per cell. **f.** CXCR2 loss reduces CXCR2 expression and PAK1 phosphorylation in RP48 liver metastases. Representative liver sections from CXCR2-WT and CXCR2-KO tumors are shown with corresponding quantification of CXCR2/DAPI and p-PAK1/DAPI signal ratios. **g.** CXCL-CXCR2 signaling increases RAC1 biosensor activity. Left: schematic of the RAC1 FRET biosensor. Right: change in Acceptor : Donor ratio in CXCR2-WT and CXCR2-KO H82 cells after vehicle or CXCL1 treatment. CXCL1 increased biosensor activity in WT but not KO cells. **h.** Doxycycline-inducible RAC1^Q61L^ increases RAC1 expression and PAK1 phosphorylation in CXCR2-KO H82 cells without increasing FAK1 phosphorylation. Representative immunoblots are shown; quantification is provided in Supplementary Figure 5d. **i.** Constitutively active RAC1 rescues migration of CXCR2-deficient non-adherent SCLC cells. Transwell migration of TetON-RAC1^Q61L^ H82 cells is shown in the absence or presence of doxycycline. **j.** Constitutively active RAC1 rescues migration of CXCR2-deficient adherent SCLC cells. Representative DMS-273 transwell images and quantification are shown in the absence or presence of doxycycline. **k.** Constitutively active RAC1 restores F-actin organization in CXCR2-KO H82 cells. Representative phalloidin staining and quantification are shown in the absence or presence of doxycycline. **l.** Constitutively active RAC1 increases SCLC-EC interaction strength in CXCR2-KO H82 cells. G-baToN flow-cytometry profiles and quantification show increased GFP transfer to HUVEC receiver cells after RAC1^Q61L^ induction. **m.** Schematic of MOBA-seq analysis of Rac1 function in SCLC metastasis. Barcoded RP48-Cas9 cells carrying three sgRac1 and three control sgRNAs were transplanted into mice and analyzed at 1 and 3 weeks. **n.** *Rac1* inactivation depletes SCLC metastatic lineages in the liver. Stacked bars show genotype contributions across individual mice and pre-transplant pools. **o.** *Rac1* inactivation reduces liver metastatic colony number. Colony-size scatter plots and marginal histograms compare sgCtrl and sgRac1 lineages. **p.** *Rac1* inactivation suppresses metastatic seeding at 1 and 3 weeks. Relative metastatic seeding is shown for individual control and Rac1-targeting sgRNAs. **q.** *Rac1* inactivation suppresses later clonal expansion. Colony-size density distributions show limited divergence at 1 week and marked suppression of sgRac1 outgrowth by 3 weeks.

To test whether direct endothelial-SCLC contact induces a small-GTPase-associated transcriptional program, we co-cultured surface GFP (sGFP)-expressing HUVECs with anti-GFP-expressing H82 cells at a 1:1 ratio for 24 h. Following G-baToN labeling, H82 cells were isolated by flow cytometry and separated into GFP-positive endothelial-contacted (TOUCH) and GFP-negative uncontacted (UNTOU) populations for bulk RNA sequencing. Transcriptomic analysis revealed distinct gene-expression programs among MONO, TOUCH, and UNTOU H82 cells. Gene-set enrichment analysis identified differential regulation of stress-response, and proliferation-associated programs, while Reactome analysis showed enrichment of Rac/Rho-family GTPase and adhesion-associated pathways in TOUCH H82 cells, including RND1 and RHOQ GTPase cycles and integrin cell-surface interactions (**Figure S4k-m**). These findings support the induction of a small-GTPase-and adhesion-associated transcriptional re-programing in SCLC following direct endothelial contact.

These data motivated us to test RAC1, a well-established regulator of actin polymerization and cell motility^39, 40^. CXCL stimulation induced time-dependent phosphorylation of FAK1 and PAK1 in SCLC cells (**Figure 5c**; **Figure S5a**). CXCR2 loss attenuated CXCL-induced FAK1 and PAK1 phosphorylation in H82 and H446 cells (**Figure 5d**; **Figure S5b**), whereas CXCR2 overexpression increased FAK1/PAK1-associated signaling (**Figure S5c**). Together, these results support CXCR2-dependent engagement of a RAC1-associated cytoskeletal signaling program.

To examine the spatial organization of this signaling response, we performed immunofluorescence staining in adherent DMS-273 cells. CXCL stimulation increased PAK1 phosphorylation with enrichment at the leading edge of migrating cells (**Figure 5e**). In liver metastases, CXCR2-KO RP48 lesions showed reduced PAK1 phosphorylation relative to CXCR2-WT lesions (**Figure 5f**). We next used a FRET-based RAC1 biosensor that reports RAC1 engagement with a PAK1-like binding domain^41^. CXCL1 increased the acceptor:donor ratio in CXCR2-WT H82 cells, whereas this response was strongly attenuated in CXCR2-KO cells (**Figure 5g**). Together, these results support a model in which CXCL-CXCR2 signaling engages RAC1-associated signaling and RAC1-dependent cytoskeletal remodeling rather than establishing RAC1 activation solely from changes in total protein abundance.

### Constitutively active RAC1 rescues CXCR2-KO phenotypes

To test whether RAC1 reactivation is sufficient to rescue CXCR2-loss phenotypes, we engineered CXCR2-KO SCLC cells to express a doxycycline (Dox)-inducible constitutively active RAC1 mutant (TetON-RAC1^Q61L^)^42^. Dox treatment increased RAC1 expression and PAK1 phosphorylation without increasing FAK1 phosphorylation in CXCR2-KO H82 cells (**Figure 5h**; **Figure S5d**). RAC1^Q61L^ induction restored migration in non-adherent H82 and adherent DMS-273 CXCR2-KO cells (**Figure 5i-j**), reinstated F-actin organization in CXCR2-KO H82 cells (**Figure 5k**), and increased SCLC-EC interaction strength in the G-baToN assay (**Figure 5l**). These rescue experiments functionally place RAC1 downstream of CXCR2 and support RAC1 as a key effector of CXCR2-dependent migratory and cytoskeletal phenotypes.

### RAC1 is essential for SCLC liver metastasis

To determine whether RAC1 is required for SCLC liver metastasis in vivo, we performed MOBA-seq in RP48 cells using three independent sgRNAs targeting Rac1 and three control sgRNAs (**Figure 5m**). At three weeks post-transplantation, Rac1-deficient lineages were depleted from liver metastases across mice and sgRNAs (**Figure 5n**; **Figure S6a**). Rac1 inactivation markedly reduced the number of liver metastatic colonies (**Figure 5o**; **Figure S6b**) and suppressed metastatic seeding at both one and three weeks (**Figure 5p**). Colony-size distributions further showed that Rac1 loss limited later clonal expansion, with a pronounced divergence from control lineages by three weeks (**Figure 5q**; **Figure S6c**). Together, these results demonstrate that Rac1 is required for both metastatic seeding and subsequent clonal outgrowth in this model.

### Targeting CXCR2-RAC1 signaling suppresses SCLC migration and SCLC-EC interactions

Our results identify CXCL-CXCR2-RAC1 signaling as an important regulator of SCLC metastatic phenotypes. We therefore asked whether pharmacologic inhibition of CXCR2 or RAC1 could suppress pro-metastatic cellular behaviors without compromising cell viability. Three human SCLC cell lines were treated with the CXCR2 inhibitor AZD5069 or the RAC1 inhibitor NSC23766 across a range of concentrations^43, 44^. Neither compound significantly affected cell viability (**Figure 6a**). We then evaluated the effect of these inhibitors on migration. Both AZD5069 and NSC23766 significantly reduced transwell migration in adherent SCLC cells (**Figure 6b-6c**).

**Figure 6:**
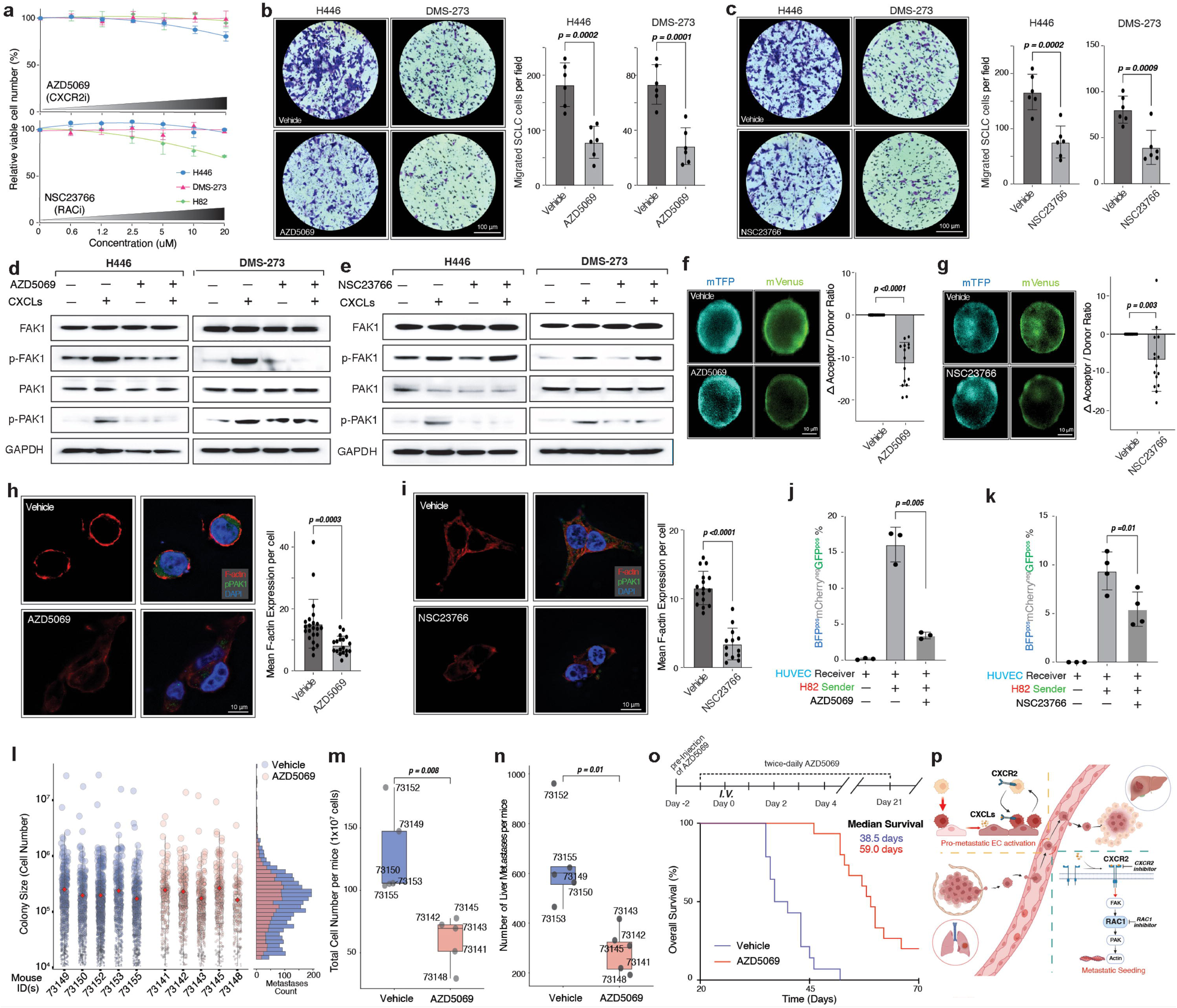
Pharmacologic CXCR2 blockade and RAC1 inhibition suppress SCLC metastatic phenotypes. **a.** AZD5069 and NSC23766 do not substantially reduce SCLC viable cell number across the indicated concentration ranges. H446, DMS-273, and H82 cells were treated with the CXCR2 inhibitor AZD5069 or the RAC1 inhibitor NSC23766 and analyzed by CCK-8 assay. **b.** AZD5069 suppresses migration of adherent SCLC cells. Representative H446 and DMS-273 transwell images and quantification are shown for vehicle and AZD5069 treatment. **c.** NSC23766 suppresses migration of adherent SCLC cells. Representative H446 and DMS-273 transwell images and quantification are shown for vehicle and NSC23766 treatment. **d.** AZD5069 attenuates CXCL-induced FAK1 and PAK1 phosphorylation. Representative immunoblots of H446 and DMS-273 cells treated with vehicle, CXCLs, AZD5069, or the combination are shown; quantification is provided in Supplementary Figure 5e. **e.** NSC23766 preferentially attenuates CXCL-induced PAK1 phosphorylation. Representative immunoblots of H446 and DMS-273 cells treated with vehicle, CXCLs, NSC23766, or the combination are shown; quantification is provided in Supplementary Figure 5f. **f.** AZD5069 reduces RAC1 FRET biosensor activity in H82 cells. Left: Representative donor (mTFP) and acceptor (mVenus) images; Right, change in acceptor:donor ratio relative to vehicle. **g.** NSC23766 reduces RAC1 FRET biosensor activity in H82 cells. Left: Representative donor and acceptor images; Right, change in acceptor:donor ratio relative to vehicle. **h.** AZD5069 reduces F-actin organization in DMS-273 cells. Representative phalloidin staining and quantification of mean F-actin signal per cell are shown. **i.** NSC23766 reduces F-actin organization in DMS-273 cells. Representative phalloidin staining and quantification of mean F-actin signal per cell are shown. **j.** AZD5069 reduces SCLC-endothelial interaction strength. G-baToN quantification shows reduced GFP transfer from H82 sender cells to HUVEC receiver cells after AZD5069 treatment. **k.** NSC23766 reduces SCLC-endothelial interaction strength. G-baToN quantification shows reduced GFP transfer from H82 sender cells to HUVEC receiver cells after NSC23766 treatment. **l.** AZD5069 suppresses RP48 SCLC liver metastatic seeding. MOBA-seq colony-size plots compare vehicle-and AZD5069-treated mice, with marginal distributions shown at right. **m.** AZD5069 reduces RP48 SCLC liver metastatic burden per mouse. **n.** AZD5069 reduces the number of RP48 SCLC liver metastatic colonies per mouse. **o.** AZD5069 pre-treatment prolongs overall survival. Mice received AZD5069 beginning before intravenous RP48 transplantation and twice-daily treatment during the indicated interval; Kaplan-Meier curves show median survival for vehicle and AZD5069 cohorts. **p.** Proposed model of reciprocal CXCL-CXCR2-RAC1 signaling during SCLC metastatic seeding. SCLC-endothelial interactions induce endothelial CXCL expression; CXCLs engage tumor-cell CXCR2 and promote RAC1-dependent cytoskeletal remodeling, migration, endothelial engagement, and metastatic establishment.

To connect these pharmacologic effects to pathway activity, we analyzed FAK1 and PAK1 signaling in H446 and DMS-273 cells treated with vehicle, CXCLs, inhibitor alone, or the combination. AZD5069 attenuated CXCL-induced FAK1 and PAK1 phosphorylation (**Figure 6d**; **Figure S5e**), whereas NSC23766 preferentially reduced PAK1 phosphorylation with less effect on FAK1 (**Figure 6e**; **Figure S5f**), consistent with RAC1 functioning downstream of FAK-associated signaling. RAC1 FRET biosensor assays further showed reduced acceptor:donor ratio after treatment with either inhibitor (**Figure 6f-g**), and both compounds decreased F-actin assembly (**Figure 6h-i**) and weakened SCLC-EC interaction strength in the G-baToN assay (**Figure 6j-k**). Together, these pharmacologic data are consistent with the genetic experiments and show that inhibition of CXCR2 or RAC1-associated signaling suppresses CXCL-driven migratory and endothelial-interaction phenotypes.

### CXCR2 blockade suppresses SCLC liver metastatic seeding

Analysis of cBioPortal lung cancer cohorts showed that activating RAC1 mutations were associated with a higher frequency of metastatic disease and shorter overall survival, whereas CXCR2 loss-of-function alterations were uncommon (**Figure S7**). To test whether pharmacologic CXCR2 inhibition could suppress SCLC liver metastatic seeding in vivo, we performed MOBA-seq in mice treated with vehicle or the CXCR2 inhibitor AZD5069. AZD5069 reduced the number of seeded metastatic colonies without significantly altering mean colony size (**Figure 6l**). Consistent with a primary effect on metastatic establishment, CXCR2 inhibition decreased both total liver metastatic burden and seeded colony number per mouse (**Figure 6m-n**). We therefore tested whether peri-seeding CXCR2 blockade could improve outcome. Recipient mice treated with AZD5069 around the time of intravenous RP48 transplantation showed significantly prolonged survival (**Figure 6o**). Together, these data demonstrate that CXCR2 inhibition limits liver metastatic seeding and prolongs overall survival in this peri-seeding treatment model.

Based on these results, we propose a reciprocal cancer-endothelial signaling model in which circulating SCLC cells induce endothelial CXCL chemokine expression, while CXCLs engage tumor-cell CXCR2 and promote RAC1-dependent cytoskeletal remodeling. This signaling program enhances SCLC migration, endothelial engagement, and metastatic seeding. Genetic or pharmacologic disruption of CXCR2, and genetic disruption of RAC1, weaken these pro-metastatic phenotypes (**Figure 6p**).

## Discussion

Metastasis depends on reciprocal signaling between disseminated cancer cells and the vascular endothelium of distant organs, yet the mechanisms governing this crosstalk remain incompletely defined. Here, we identify an SCLC-induced endothelial CXCL program and show that CXCL-CXCR2 signaling promotes tumor-cell migration, endothelial engagement, and metastatic seeding. MOBA-seq quantitatively resolved the contribution of CXCR2 to metastatic establishment and subsequent clonal behavior, while transcriptomic, biosensor, rescue, and genetic experiments supported RAC1-dependent cytoskeletal remodeling downstream of CXCR2. Pharmacologic CXCR2 blockade reduced liver metastatic seeding and prolonged survival in a peri-seeding treatment model. Together, these findings define a reciprocal cancer-endothelial signaling axis that links endothelial CXCL induction to tumor-intrinsic CXCR2-RAC1 activity during metastatic establishment.

Metastatic colonization requires disseminated cancer cells to engage and adapt to the local endothelium within target organs. Our data support a reciprocal model of cancer-endothelial communication in SCLC liver metastasis in which SCLC exposure induces endothelial CXCL chemokine production, while CXCL signaling strengthens tumor migration and endothelial engagement. The upstream signals that induce endothelial CXCL expression remain to be defined. Prior work has shown that inflammatory cytokines, including IL-1 and TNF-alpha, can induce CXCL transcription in multiple cell types, including endothelial cells^45, 46^. Consistent with inflammatory pathway activation, our scRNA-seq analysis identified an SCLC-induced LSEC population enriched for TNF-associated genes. Direct cell-cell contact may also contribute to endothelial activation. Although G-baToN assays showed that CXCL signaling can strengthen SCLC-endothelial interactions in vitro, we have not directly shown that this strengthened contact further increases endothelial CXCL output. Thus, the present data support reciprocal or feed-forward signaling rather than a fully closed positive-feedback loop. Defining the tumor-derived soluble and contact-dependent signals that regulate endothelial CXCL expression will require targeted perturbation coupled to endothelial transcriptional and signaling readouts in vitro and in vivo.

Several additional questions must be addressed to build a more comprehensive model of endothelial reprogramming in SCLC metastasis. First, it remains unclear whether SCLC-induced CXCL expression is restricted to resident endothelial cells in specific metastatic target organs or instead represents a broader endothelial response that is quantitatively amplified in selected vascular beds. Our data suggest that liver endothelium is particularly responsive in vivo, whereas HUVECs can also mount a CXCL response under high-exposure co-culture conditions, supporting the possibility that both intrinsic vascular-bed identity and CTC accessibility contribute. Systematic comparisons across organ-specific endothelial populations, ideally using matched in vitro and in vivo transcriptomic profiles, will be needed for future studies. Second, the functional consequences of CXCL-high endothelial states likely extend beyond CTC chemotaxis and may involve autocrine and paracrine endothelial programs that reshape the metastatic niche. How CXCL induction intersects with vascular permeability, immune recruitment, endothelium remodeling, and angiogenic signaling, as well as which of these downstream programs are necessary for metastatic seeding, persistence, and clonal outgrowth, remain open questions for future studies. Third, our data suggest that CXCL-independent pathways may also contribute to a pro-metastatic liver endothelial response. Prior studies have implicated CSF2 in promoting breast cancer metastasis, and our scRNA-seq analysis revealed marked upregulation of CSF2 within the cytokine-releasing LSEC cluster^47^. Whether endothelial CSF2 acts in parallel with the CXCL-CXCR2 axis, converges on shared downstream programs, or promotes metastasis through an independent mechanism remains to be determined. Finally, existing strategies to target tumor vasculature have focused on anti-angiogenesis, such as targeting VEGF/VEGFR signaling. Clinical outcomes have been mixed, with benefits in some contexts but variable efficacy and toxicities, including effects on vascular integrity that could influence metastasis^48^. Our finding that liver endothelium can function as a pro-metastatic niche suggests that directly targeting cancer-endothelial communication, such as anti-CXCL neutralizing antibodies or CXCR2 antagonists, may offer new avenues for limiting metastatic establishment in SCLC. Defining the therapeutic window, optimizing dosing and timing, and evaluating efficacy in single-agent versus combination therapies, will be important priorities for future translational studies.

How metastatic cancer cells translate stromal inputs into the cellular programs that drive metastatic progression remains incompletely understood. Our study identifies the CXCR2-RAC1 axis as a key effector of CXCL-induced signaling in SCLC-EC crosstalk. However, the mechanisms by which CXCR2-RAC1 signaling reinforces cancer-endothelial adhesion remain to be defined. Cytoskeletal remodeling is a hallmark of the organization of adhesion machinery at cell-cell interfaces^49^. Our data suggest that CXCL chemokines induce CXCR2-dependent PAK1 phosphorylation that is enriched at the leading edge of migrating SCLC cells. In addition, studies in osteosarcoma have implicated the role of CXCR2 in regulating expression of the adhesion molecule VCAM1^21^. These observations support the hypothesis that CXCR2-RAC1 signaling may promote the expression and reorganization of adhesion molecules that directly strengthen SCLC binding to the endothelium. Systematic perturbation of candidate adhesion proteins downstream of CXCR2 and RAC1, coupled with quantitative in vitro and in vivo assays of cancer-endothelial interactions will be required for future research.

MOBA-seq enables quantitative dissection of genotype-specific metastatic phenotypes at single-colony resolution with near single-cell accuracy^33^. In our analyses, CXCR2 loss increased the fraction of small, non-expanding liver colonies that are consistent with a dormant-like state, but colony size alone does not establish biological dormancy. Orthogonal measurements of proliferation and cell-cycle state will be needed to determine whether CXCR2 directly controls entry into or exit from dormancy. Prior studies suggest that RAC1 activity can support cell-cycle progression^50–52^, raising the possibility that CXCR2-RAC1 signaling could influence the transition from persistence to clonal outgrowth. Immune surveillance may also contribute to these dynamics, particularly because the MOBA-seq experiments were performed in recipient mice with distinct immune backgrounds. A second notable finding is that CXCR2 and RAC1 perturbation produced different metastatic phenotypes: CXCR2 loss predominantly impaired seeding, whereas Rac1 loss suppressed both seeding and subsequent clonal expansion. This divergence is consistent with RAC1 functioning as a broader signaling node that integrates multiple upstream inputs, including growth-factor pathways such as PDGF and EGF^53, 54^. Defining the upstream regulators that converge on RAC1 in vivo will be important for determining whether RAC1 can be safely and selectively targeted in established metastatic disease.

Standard treatment for limited-stage SCLC includes platinum-based chemotherapy and thoracic radiation, with immune-based consolidation used in selected settings^55^. Relapse remains common, underscoring the need for strategies that limit metastatic establishment as well as established tumor burden. Our MOBA-seq analyses show that CXCR2 inactivation reduces metastatic seeding and increases the fraction of small, non-expanding liver colonies, while our in vivo pharmacology shows that peri-seeding AZD5069 treatment reduces liver metastatic burden and prolongs survival. These findings support evaluation of CXCR2 blockade as a metastasis-prevention strategy in settings where dissemination risk is high, while recognizing that efficacy against established metastatic lesions was not tested here. RAC1 perturbation had broader effects on both seeding and clonal expansion, but RAC1 inhibition was not evaluated in vivo and NSC23766 has known off-target pharmacology. Future studies should therefore prioritize genetically and pharmacologically selective RAC1 perturbation, define dosing windows, and test whether CXCR2 or RAC1 targeting can complement standard therapies without compromising vascular or immune function.

In summary, our study identifies CXCL-CXCR2 signaling as a mediator of cancer-endothelial interactions during early SCLC metastatic seeding. We show that endothelial CXCL induction is coupled to tumor-intrinsic CXCR2 signaling and RAC1-dependent cytoskeletal remodeling, and that CXCR2 blockade limits metastatic establishment in vivo. These findings provide a mechanistic framework for how endothelial stromal inputs are translated into tumor-cell programs that support metastatic colonization and identify the CXCL-CXCR2-RAC1 axis as a potential therapeutic vulnerability in metastatic SCLC.

## Supporting information

Suppemental Figure 1

Suppemental Figure 2

Suppemental Figure 3

Suppemental Figure 4

Suppemental Figure 5

Suppemental Figure 6

Suppemental Figure 7

## CONFLICT OF INTERESTS

The authors declare no potential conflicts of interest

## ACKNOWLEDGMENTS

We dedicate this work to the memory of Gui Zhang, an exceptional collaborator and dear friend who helped build the technical foundation of this study for the Tang lab. We thank the staff at Division of Comparative Medicine (DCM) at Washington University for expert animal care, Anatomic and Molecular Pathology (AMP) Core Labs, Siteman Flow Cytometry Core (SFC), and the WashU Genome Technology Access Center (GTAC) for experimental support. We thank Kate Sutherland for sharing RP48 and RP116 cells. We thank Dr. Brett Herzog for helpful comments. This work was supported by NIH R00-CA256039 (to R.T.), Cancer Research Foundation Young Investigator Award (to R.T.), the Phi Beta Psi Sorority Research Grant (to R.T.), the American Lung Association Innovation Award (to R.T.), U01CA294532 (to L.D., F.C), U54AG075934 (to L.D., F.C), and R01CA260112 (to L.D., F.C). The funders had no role in study design, data collection and analysis, and decision to publish or preparation of the manuscript.

## AUTHOR CONTRIBUTIONS

Z.Y. and R.T. conceived the project and designed the experiments. Z.Y. led experimental data production with contributions from all authors. A.X. and R.T. led the data analysis. R.T. oversaw the project. A.X. and R.T. wrote the manuscript with input from all authors.

## Figure Legends

**Supplementary Figure 1: Transcriptomic characterization of SCLC-induced endothelial states and human metastatic endothelial subtypes. a.** Reference UMAP from the MacParland human liver single-cell dataset (GSE115469) used for endothelial subtype annotation and validation.

**b.** Projection of label-transfer prediction scores onto the reference UMAP for mapped LSEC populations.

**c.** Cell-type composition across mapped samples, showing the relative proportions of periportal LSECs, central venous LSECs, with little hepatocyte contaminations.

**d.** Differential-expression volcano plots comparing EC2-EC5 with EC1 in the LSEC scRNA-seq dataset.

**e.** Violin plots of CXCL1, CXCL2, CXCL3, and CXCL8 expression across EC1-EC5, highlighting the CXCL-high EC4 state.

**f.** PCA of bulk RNA-seq profiles from HUVEC monoculture and H82-HUVEC co-culture samples.

**g.** Heatmap of top differentially expressed genes between HUVEC monoculture and H82-HUVEC co-culture conditions.

**h.** Gene-set enrichment analysis of HUVEC monoculture versus H82-HUVEC co-culture conditions.

**i.** Cnetplot showing core enriched genes contributing to co-culture-associated pathway changes in HUVECs.

**j.** Endothelial cell subtype composition in colorectal cancer liver and lung metastases, showing LSECs as the predominant liver parenchymal endothelial population and alveolar ECs as a major lung parenchymal endothelial population.

**k.** Representative marker genes used to annotate endothelial subtypes in the liver and lung metastatic datasets.

**Supplementary Figure 2: Conservation and validation of SCLC-induced endothelial CXCL responses in mouse and human models. a.** Phylogenetic analysis of CXCL chemokine sequences across mouse and human, showing conservation of the CXCR2 ligand family.

**b.** Schematic of primary mouse LSEC isolation and co-culture with KP-1 SCLC cells before endothelial re-isolation and RT-qPCR analysis.

**c.** RT-qPCR analysis of endothelial transcripts in monocultured versus KP-1-co-cultured mouse LSECs. Data are shown as relative mRNA fold change.

**d.** H82 SCLC co-culture increases CXCL1 protein expression in HUVECs. Representative CXCL1/CD31/DAPI immunofluorescence images and quantification are shown for monoculture and co-culture conditions.

**e.** RP48 SCLC transplantation preferentially increases endothelial CXCL1 signal in the liver. Representative CXCL1/CD31/DAPI immunofluorescence images from brain, lung, and liver of sham-treated or RP48-transplanted mice are shown with tissue-specific quantification.

**Supplementary Figure 3: CXCR2 inactivation differentially affects metastatic colony-size and outgrowth metrics. a.** Cxcr2 inactivation alters the relative 90th-percentile metastatic colony size across liver and lung tissues and time points in RP48 SCLC.

**b.** Colony-size density distributions for sgCtrl and sgCxcr2 RP48 metastases in liver and lung across selected time points.

**c.** Relative 90th-percentile liver metastatic colony size for control and CXCR2-, CXCR4-, or NFIB-targeted H82 SCLC lineages at 3 weeks.

**d.** Liver colony-size distributions for control, CXCR2-, CXCR4-, and NFIB-targeted H82 SCLC lineages at 3 weeks.

**e.** Relative peak-mode colony-size metric for the indicated H82 genotypes.

**f.** Gaussian mixture modeling of H82 sgCtrl and sgCXCR2 liver colony-size distributions, showing genotype-associated differences in the inferred dormant-like, non-expanding colony fraction.

**Supplementary Figure 4: CXCR2 signaling regulates SCLC transcriptional states and Rac/Rho-associated pathways. a.** PCA of H82 transcriptomes across CXCR2-WT, CXCR2-KO, CXCR2-overexpressing, and CXCL-treated conditions.

**b.** Volcano plot of differentially expressed genes between CXCR2-KO and CXCR2-WT H82 cells.

**c.** Heatmap of differentially expressed genes between CXCR2-KO and CXCR2-WT H82 cells.

**d.** Pathway-network analysis of programs altered by CXCR2 loss, highlighting cytoskeletal, adhesion, Rap1, Ras, and related signaling pathways.

**e.** Volcano plot of differentially expressed genes in CXCL-stimulated versus control H82 cells.

**f.** Heatmap of differentially expressed genes in CXCL-stimulated versus control H82 cells.

**g.** Volcano plot of differentially expressed genes in CXCR2-overexpressing versus CXCR2-WT H82 cells.

**h.** Heatmap of differentially expressed genes in CXCR2-overexpressing versus CXCR2-WT H82 cells.

**i.** GSEA summary of Rac/Rho axis and cytoskeletal pathways altered by CXCR2 perturbation.

**j.** Metastatic-seeding effects of Rac-, Rap-, and Rho-family gene perturbations across liver, lung, and brain in MOBA500 datasets^33^.

**k.** Principal component analysis of transcriptomic profiles from anti-GFP-expressing H82 cells cultured alone (MONO) or co-cultured with surface-GFP–expressing HUVECs. Following G-baToN labeling, H82 cells were sorted into GFP-positive endothelial-contacted (TOUCH) and GFP-negative uncontacted (UNTOU) populations for RNA sequencing.

**l.** Gene-set enrichment analysis comparing transcriptional programs among MONO, TOUCH, and UNTOU H82 cells, highlighting differential enrichment of stress-response and proliferation-associated pathways.

**m.** Reactome analysis showing enrichment of Rac/Rho-family GTPase and adhesion-associated pathways in endothelial-contacted TOUCH H82 cells, including RND1 and RHOQ GTPase cycles and integrin cell-surface interactions.

**Supplementary Figure 5: Quantification of FAK1/PAK1 signaling changes associated with CXCL-CXCR2-RAC1 perturbation. a.** Quantification of p-FAK1/FAK1 and p-PAK1/PAK1 ratios from CXCL time-course immunoblots in H82 and H446 cells corresponding to Figure 5c.

**b.** Quantification of p-FAK1/FAK1 and p-PAK1/PAK1 ratios in CXCR2-WT and CXCR2-KO H82 and H446 cells with or without CXCL treatment corresponding to Figure 5d.

**c.** Representative immunoblots and quantification showing increased FAK1/PAK1-associated signaling in CXCR2-overexpressing H82 and H446 cells.

**d.** Quantification of FAK1 and PAK1 phosphorylation after doxycycline-induced RAC1Q61L expression in CXCR2-KO H82 cells corresponding to Figure 5h.

**e.** Quantification of FAK1 and PAK1 phosphorylation in H446 and DMS-273 cells treated with CXCLs and/or AZD5069 corresponding to Figure 6d.

**f.** Quantification of FAK1 and PAK1 phosphorylation in H446 and DMS-273 cells treated with CXCLs and/or NSC23766 corresponding to Figure 6e.

**Supplementary Figure 6: MOBA-seq analysis of Rac1 function in SCLC liver metastasis in C57BL/6 mice. a.** Genotype composition of sgCtrl and sgRac1 barcoded RP48 lineages across pre-transplant pools and individual mouse livers.

**b.** Colony-size scatter plots and marginal distributions comparing sgCtrl and sgRac1 liver metastatic lineages.

**c.** Colony-size density distributions for sgCtrl and sgRac1 lineages at 1 and 3 weeks, illustrating the later effect of Rac1 loss on clonal expansion.

**Supplementary Figure 7: CXCR2 and RAC1 genomic alterations are associated with metastatic status and survival in human lung cancer datasets. a.** Distribution of TNM metastatic status in patients with CXCR2 wild-type tumors versus tumors carrying CXCR2 loss-of-function alterations. Bar plots show the proportions of M0 and M1 cases.

**b.** Distribution of TNM metastatic status in patients with RAC1 wild-type tumors, RAC1 loss-of-function alterations, or activating oncogenic RAC1 mutations. Bar plots show the proportions of M0 and M1 cases.

**c.** Kaplan-Meier analysis of overall survival comparing patients with activating RAC1 mutations versus RAC1 wild-type tumors.

## Methods

### Mouse research

The use of mice for the current study was approved by the Institutional Animal Care and Use Committee at Washington University, protocol no. 23-0283. We used mice of both sexes, 8-15 weeks of age at the time of tumor initiation. Mice were housed at NRB Basement barrier facility under a 12-hour light/12-hour dark cycle with dark hours from 18:30 to 06:30. Mice were housed at 68-73 °F and 40-60% relative humidity.

### Cells, Plasmids and Reagents

HEK-293, H82, H446, DMS-273, and HUVEC cells were purchased from ATCC; KP-1 cells were kindly provided by Julien Sage (Stanford School of Medicine); RP48 and RP116 small cell lung cancer cells were generated in the Kate Sutherland laboratory; primary human liver sinusoidal endothelial cells were obtained from ScienCell Research Laboratories (catalog no. 5000); HEK-293 cells were cultured in DMEM containing 10% FBS, 100 units/mL penicillin and 100 μg/mL streptomycin. DMS-273, H446 and H82 cells were cultured in RPMI1640 medium containing 10% FBS, 100 units/mL penicillin and 100 μg/mL streptomycin. RP48 and RP116 cells were cultured in DMEM/F12 (Thermo Fisher), 10% (vol/vol) FBS, 6 ng/mL Mouse EGF (FUJIFILM IRVINE SCIENTIFIC INC), and 4 µg/mL Hydrocortisone (Cayman Chemical). HUVECs were cultured in Vascular Cell Basal Medium (ATCC, PCS-100-030) with Endothelial Cell Growth Kit (ATCC, PCS-100-041); All cell lines were confirmed to be mycoplasma negative (MycoAlert Detection Kit, Lonza). All plasmids used in this study are available from our laboratory upon request.

Primary antibodies included anti-CXCL1 (PeproTech, 500-P115-50UG); anti-CD31 (Abcam, clone RM0032-1D12, ab56299); anti-CXCR2 (Antibodies.com, A9177); anti-FAK (Cell Signaling Technology, clone D2R2E, 13009S); anti-phospho-FAK (Tyr397; Cell Signaling Technology, clone D20B1, 8556S); anti-RAC1/CDC42 (Cell Signaling Technology, 4651S); anti-PAK1/2/3 (Cell Signaling Technology, 2604S); anti-phospho-PAK1 (Thr423)/PAK2 (Thr402) (Cell Signaling Technology, 2601S); and anti-GAPDH (Cell Signaling Technology, clone D16H11, 5174S).

### Lentiviral Vector Packaging

Lentiviral vectors were produced using polyethylenimine (PEI)-based transfection of 293T cells with the plasmids indicated in Supplementary File 1, along with delta8.2 and VSV-G packaging plasmids in 150-mm cell culture plates. Sodium butyrate (Sigma-Aldrich, B5887) was added 8 hours after transfection to achieve a final concentration of 20 mM. Medium was refreshed 24 hours after transfection. 20 mL of virus-containing supernatant was collected 36, 48, and 60 hours after transfection. The three collections were then pooled and concentrated by ultracentrifugation (25,000 rpm for 1.5 hours), resuspended overnight in 100 µL PBS, then frozen at -80°C.

### Generation of Stable Cell Lines

Parental cells were seeded at 50% confluency in a 6-well plate the day before transduction (day 0). The cell culture medium was replaced with 2 mL fresh medium containing 8 µg/mL hexadimethrine bromide (Sigma-Aldrich, H9268-5G), 20 µL ViralPlus Transduction Enhancer (Applied Biological Materials Inc., G698) and 40 µL concentrated lentivirus and cultured overnight (Day 1). The medium was then replaced with complete medium and cultured for another 24 hours (Day 2). Cells were transferred into a 100 mm cell culture dish with appropriate amounts of puromycin (Dose used: 293T: 2 µg/mL) and selected for 48 hours (Day 3).

To generate CXCR2-knockout cells by CRISPR/Cas9 editing, human and mouse Cas9+ SCLC cell lines were transduced with lentiviral sgRNAs targeting species-appropriate CXCR2/Cxcr2 sequences. After 3 days of puromycin selection, 100,000 cells from each cell line were collected for DNA extraction using the QIAGEN DNeasy Blood and Tissue Kit (Cat# 69506). DNA was quantified by NanoDrop, and 200 ng was used for PCR amplification of human and mouse CXCR2/Cxcr2. Amplicons were purified using the QIAGEN QIAquick PCR Purification Kit (Cat# 28106) and analyzed on a 1% agarose gel for size verification (expected sizes: human, 344 bp; mouse, 255 bp). Samples were submitted to Azenta for Sanger sequencing, and traces were analyzed using Synthego ICE to estimate CRISPR editing efficiency.

### Transwell Co-culture Assay

The Corning® Transwell® polycarbonate membrane cell culture inserts were purchased from Corning Inc (3422: CS, Corning, NY). SCLC cells were seeded in the upper chamber inserts of the transwell (1×10^5^/insert). The inserts were then put back into the receiver plate filled with SCLC culture medium plus CXCL1/2/3 cocktail and incubated at 37 °C in a humidified 5% CO2 incubator for 24-48 hrs. After incubation, the transwell inserts were washed with PBS three times and fixed with 4% paraformaldehyde for 10 min at room temperature, followed by washing with PBS. Cells were stained with Wright-Giemsa for acquiring images or DAPI for cell counting.

### Cell viability assay

For cell proliferation (CCK-8) assays, cells were seeded in 96-well plates at a density of 5,000 cells per well and allowed to adhere overnight in regular growth medium (day 0). Cells were then cultured under the conditions indicated in each figure for 72 hours. Relative viable cell number was measured using Cell Counting Kit-8 (Bimake, B34304) according to the manufacturer’s instructions.

### Western Blot

5×106 SCLC cells were cultured in a 100-mm cell culture dish for 24 hours. Cells were lysed in RIPA buffer (50 mM Tris-HCl (pH 7.4), 150 mM NaCl, 1% Nonidet P-40, and 0.1% SDS) and incubated at 4 °C with continuous rotation for 30 minutes, followed by centrifugation at 12,000 × rcf for 10 minutes. The supernatant was collected, and the protein concentration was determined by BCA assay (Thermo Fisher Scientific, 23250). Protein extracts (20–50 μg) were resolved by 10% SDS-PAGE and transferred onto PVDF membranes. The membranes were blocked with 5% non-fat milk in TBS with 0.1% Tween 20 (TBST) at room temperature for one hour, followed by incubation with primary antibodies diluted in TBST at 4 °C overnight. After three 10-min washes with TBST, the membranes were incubated with the appropriate secondary antibody conjugated to HRP diluted in TBST (1:10000) at room temperature for 1 hour. After three 10-minutes washes with TBST, protein expression was quantified with enhanced chemiluminescence reagents (Fisher Scientific, PI80196).

### Immunofluorescent staining

Cells were seeded on cover glasses placed in a 24-well plate and allowed to adhere overnight. Cells were then treated with CXCL1/2/3 cocktail for 24 h, fixed with 1% PFA in PBS, permeabilized with 0.1% Triton X-100/PBS for 5 min, followed by blocking with 10% goat serum and incubation with Texas Red®-X phalloidin or p-PAK1 antibody at 4 °C overnight. FITC-conjugated secondary antibodies were used to detect target protein expression.

### Flow Cytometry

Metastasis-bearing mouse tissues were dissected, cut into small pieces and digested in 5 mL tissue digestion medium (3.5 mL HBSS-Ca2+-free, 0.5 mL Trypsin-EDTA [0.25%], 5 mg Collagenase IV [Worthington], 25 U Dispase [Corning]) for 30 min in hybridization chamber at 37°C with rotation. Digestion is then neutralized by adding 5 mL ice cold Quench Solution (4.5 mL L15 media, 0.5 mL FBS, 94 µg DNase). Single-cell suspensions were generated by filtering through a 40-µM cell strainer, spinning down at 500 rcf for 5 min and washed with PBS twice. Digested single-cell suspensions were subjected to a 30% Percoll gradient for separation of cancer and immune cells from hepatocytes and fat. In short, a stock isotonic Percoll (SIP) solution was made combining one-part 10X HBSS (14065056, ThermoFisher Scientific) with nine-parts of Percoll (P1644, Millipore Sigma). Digested liver samples were centrifuged at 300 x g for 8 minutes to pellet. Supernatant was removed, and samples were resuspended in 100uL of Cell Staining Buffer (420201, BioLegend) containing LIVE DEAD Blue staining (1:400 dilution) and incubated at room temperature (RT) in the dark for 10 minutes. Samples were then washed and resuspended with the surface antibody cocktail prepared in a 1:1 mixture of Cell Staining Buffer and BD Horizon Brilliant Stain buffer (566349, BD Biosciences) for 25 minutes at room temperature in the dark. After staining, samples were washed once with Cell Staining Buffer and resuspended in 100uL buffer prior to acquiring data on a Cytek Aurora Spectral Flow Cytometer. Analysis of acquired data was performed using FlowJo_v.10.10.0.

### G-baToN assay

Cell-cell interactions were quantified using the GFP-based Touching Nexus (G-baToN) system, as previously described^30^. For SCLC-endothelial interaction assays, mCherry-labeled H82 cells expressing surface GFP (sGFP) were used as sender cells, and BFP-labeled HUVECs expressing a cell-surface anti-GFP nanobody (αGFP) were used as receiver cells. Sender and receiver cells were mixed and co-cultured for 24 h under orbital shaking in the presence of the indicated treatment. Following co-culture, cells were collected, dissociated into single-cell suspensions, and analyzed by flow cytometry. Interaction-dependent labeling was quantified as the percentage of GFP+ cells within the viable receiver population.

For non-endothelial receiver assays, H82 cells expressing surface mCherry were used as sender cells, and anti-mCherry-expressing HEK-293 cells carrying GFP control, CXCL1-GFP, or CXCL2-GFP constructs were used as receiver cells. Sender and receiver cells were co-cultured for 24 h under orbital shaking. Following flow-cytometric analysis, HEK-293T receiver cells were identified by GFP expression, and interaction-dependent labeling was quantified as the percentage of mCherry+ cells within the viable GFP+ receiver-cell population.

### RAC1 FRET reporter assay

RAC1 activity was measured using a genetically encoded mTFP-mVenus Förster resonance energy transfer (FRET) biosensor, as previously described^41^. The biosensor contains full-length RAC1 and a PAK1-derived RAC1-binding domain positioned between the donor and acceptor fluorescent proteins. CXCR2-WT and CXCR2-KO H82 cells engineered to express the RAC1 FRET biosensor were plated in glass-bottom dishes and imaged before and after treatment with vehicle, CXCL1, AZD5069, or NSC23766. Donor and acceptor emission images were acquired and RAC1 activity is quantified by the change in acceptor:donor ratio. Mean mTFP donor and mVenus acceptor fluorescence intensities were quantified within each cell. RAC1 biosensor activity was expressed as the acceptor:donor fluorescence ratio. For each cell, the change in acceptor:donor ratio (ΔA:D) was calculated relative to its pretreatment baseline:

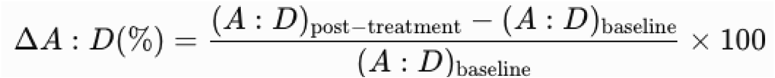

### 3D microfluidic cancer cell-endothelial co-culture

The master mold of microfluidic chips was fabricated using a 3D printer (Titan HD, Kudo3D Inc. Dublin, CA). The surface of the molds was spray-coated with silicone mold release (CRC, cat. No.: 03300) and PDMS (polydimethylsiloxane, Sylgard 182, Dow Corning) was poured on it. After heat curing at 65°C for approximately 5 hours, the solidified PDMS replica was peeled off from the mold. Holes were made at both ends of each channel in the PDMS replica using a biopsy punch. The PDMS replica was then bonded to precleaned microscope glass slides (Fisher Scientific) through plasma treatment (Harrick Plasma PDC-32G, Ithaca, NY). Microfluidic chips were UV-treated overnight for sterilization before cell seeding. A basement membrane extract (BME) hydrogel (Cultrex™ reduced growth factor basement membrane matrix type R1, Trevigen, Cat #: 3433-001-R1) was injected into the middle hydrogel channel of the chips placed on a cold pack and then transferred to rectangular 4-well cell culture plates (Thermo Scientific, Cat #: 267061) followed by incubation at 37°C in a cell culture incubator for 30 minutes for gelation. After gelation, 10 uL of human umbilical vein endothelial cells (HUVECs) resuspended at the density of ∼ 1 × 106 cells/mL was injected to the blood channel of the chips and endothelial cell growth medium wa added to the other side channel. After incubation for 3h for cells to adhere, old medium in both side channels was replaced with fresh medium. The next day, samples were placed on a rocking see-saw shaker (OrganoFlow® L, Mimetas) that generates a pulsatile bidirectional flow to mimic the dynamic native environment and cultured for 4 more days to form a complete endothelium. Cell culture medium was changed every other day.

### scRNA-seq data analysis

Raw FASTQ files from five scRNA-seq samples were processed with CellRanger v8 (10X Genomics) for alignment to the human reference genome (GRCh38-2024-A) and generation of feature barcode matrices. Matrices were imported to Seurat v5 individually and merged with sample-specific cell barcode prefixes. Samples were annotated by experimental condition: Monoculture (B1), Coculture-1 (B2, B3), and Coculture-2 (B4, B5).

Quality control filtering retained cells with ≥2,000 UMIs, ≥1,000 detected genes, log10(genes per UMI) >0.8, and <20% mitochondrial content. Genes expressed in fewer than 10 cells were excluded from downstream analyses. Normalization was performed using SCTransform with regression of mitochondrial gene percentage. Principal component analysis was conducted, followed by shared nearest neighbor graph construction (FindNeighbors, dims = 1:40) and clustering using the smart local moving (SLM) algorithm (FindClusters, algorithm = 3, resolution = 0.39). Uniform Manifold Approximation and Projection (UMAP) was used for dimensionality reduction and visualization. Cell types were annotated using SingleR with the Human Primary Cell Atlas reference. Endothelial cells were subset based on SingleR labels, re-normalized with SCTransform, and re-clustered. Small clusters lacking distinct marker gene expression were removed, and the remaining cells were re-clustered to define final populations.

### scRNA-seq differential expression and velocity analysis

Cluster marker genes were identified using Presto with Wilcoxon rank-sum test. Pairwise differential expression analyses between clusters were performed without a log fold-change threshold. Gene set enrichment analysis was conducted using fgsea with MSigDB C7 immunologic gene sets, ranking gene by area under the curve (AUC). Single-sample enrichment scores were calculated using escape with th UCell method and MSigDB Hallmark gene sets.

Spliced and unspliced transcript counts were quantified from BAM files using velocyto to generate loom files for each sample. Loom files were loaded into Python and filtered to retain only cells present in the final Seurat object. Cell identities and UMAP coordinates were transferred from the final Seurat object to maintain consistency between analyses.

RNA velocity was estimated using scVelo with stochastic mode. Genes with ≥20 shared counts across cells were retained, and the analysis was restricted to the top 2,000 highly variable genes. First- and second-order moments were computed using 30 principal components and 30 nearest neighbors. Velocity vectors were calculated and projected onto the UMAP embedding for visualization as streamline plots. Velocity analysis was performed on the merged dataset and separately for each experimental condition (Monoculture, Coculture-1, Coculture-2).

### Xenium data analysis

Following the optimized Xenium analysis pipeline, we preprocessed and clustered Xenium data as follows: 1) After formatting each raw data as an Anndata structure, we then concatenated both groups into one Anndata with their group identifications. 2) Then, we filtered out the cells with a minimum of 40 transcripts and a minimum of 15 gene types. After that, we performed normalization first with CP10K and then with log1p. Then, we selected the top 3000 highly variable genes from a total of 5106 genes for each group. 3) We then used PCA to get the top 50 principal components, and performed neighbor analysis to find 30 closest neighbors in the feature matrix for each cell and construct a corresponding neighborhood graph. 4) Finally, we performed the Leiden clustering method to classify all cells into 15 clusters, and also performed UTAG and gene ranking for characterizing these clusters.

For Cell annotation, we annotated all cell clusters for their cell types based on the top 30 genes of each cluster and related marker genes of targeted cell types. The key cell types, like tumor cells and endothelial cells, were further confirmed by their marker genes.

### SCLC transplantation via tail vein injection

For intravenous transplantations, 2.5×10^6^ SCLC cells were injected into one of the lateral tail veins. For longitudinal Cxcr2 MOBA-seq experiments, tissues were collected at 2 days and 1, 2, and 3 weeks after transplantation. For Rac1 MOBA-seq experiments, livers were collected at 1 and 3 weeks. For HDI experiments, livers were collected 2 weeks after transplantation. Survival cohorts were monitored until predefined humane endpoints. Lung, liver, brain, bone marrow, and blood cells were collected for gDNA extraction. For paired subcutaneous transplantation experiments, 2.5 × 10 GFP-labeled CXCR2-WT and 2.5 × 10 mCherry-labeled CXCR2-KO RP48 cells were resuspended separately in Matrigel and injected into opposite flanks of the same mouse. Mice were sacrificed 28 days post-injection. Both primary SubQ tumors and metastasis-bearing liver were dissected and the weight, height, width, and length, of each tumor was measured. Maximal tumor size/burden permitted by Institutional Animal Care and Use Committee (IACUC) is 1.75 cm^3^, the maximal tumor size/burden was not exceeded in our study. IACUC approved all animal studies and procedures.

### MOBA-seq library generation

Genomic DNA was isolated from bulk tumor-bearing tissues from each mouse as previously described^33^. Briefly, benchmark monoclonal control cell lines were generated from B16F10 cells transduced by a barcoded Lenti-sgDummy/PuroR vector (sgDummy: a control sgRNA with dual Bsmb1 cloning sites, AGAGACGCTCGAGCGTCTCT) and selected for monoclonal culture. Eight to ten benchmark control cell lines with varying number of cells were added to each mouse tissue sample prior to lysis to enable the calculation of the absolute number of cancer cells in each tumor from the number of sgID-BC reads. Following homogenization and overnight proteinase K digestion, genomic DNA was extracted from the tissue lysates using standard phenol-chloroform and ethanol precipitation methods. Subsequently, Q5 High-Fidelity 2x Master Mix (New England Biolabs, M0494X) was used to amplify the sgID-BC region from 40 μg of genomic DNA in a total reaction volume of 400 μl per sample for 1st round PCR amplification. The primers used were Forward: ATA ATG TGT GTG GTA CAA AAG GTC and Reverse: GAA TTC CAT GTT AAT TAA GCC ATA GGC. The PCR products were purified with the QIAquick PCR Purification Kit (Qiagen, 28104). Q5 High-Fidelity 2x Master Mix (New England Biolabs, M0494X) was used to amplify the sgID-BC region from 10% of the purified PCR product in a total reaction volume of 200 μl per sample for the NGS PCR amplification. The unique dual-indexed primers used were Forward: AAT GAT ACG GCG ACC ACC GAG ATC TAC AC-8 nucleotides for i5 index-ACA CTC TTT CCC TAC ACG ACG CTC TTC CGA TCT-4 to 7 random nucleotides for increased diversity-AGG GTT ACA GTT TAG TCA CCA TA and Reverse: CAA GCA GAA GAC GGC ATA CGA GAT-6 nucleotides for i7 index-GTG ACT GGA GTT CAG ACG TGT GCT CTT CCG ATC T-9 to 6 random nucleotides for increased diversity-CGA CTC GGT GCC ACT TTT TC. The PCR products were purified with the QIAquick PCR Purification Kit (Qiagen, 28104). The concentration and quality of the purified libraries were determined using the Invitrogen 1X dsDNA High Sensitivity kit (Invitrogen, Q33231) on the Invitrogen Qubit 4 Fluorometer (Invitrogen, Q33238). The libraries were assigned one to ten million reads based on tissue type to ensure adequate sequencing coverage, purified with the QIAquick Gel Extraction kit (Qiagen, 28704), and sequenced (read length 2×150bp) on the NovaSeq X Plus platform (GTAC@MGI, Novogene).

### Processing of paired-end reads to identify the sgID and barcode

Sequencing of MOBA-seq libraries produces reads that are expected to contain a 23-nucleotide barcode (BC) of the form GTTNNNNNNNNNNNNNNNNNATG, where each of the 17 Ns represents a random nucleotide, followed by a 17 to 21 nucleotide sgID. Each sgID has a one-to-one correspondence with an sgRNA in the lentiviral-sgRNA library (sgID dictionary). Note that all sgID sequences in the viral pool differ from each other by at least five nucleotides such that incorrect sgID assignment due to PCR or sequencing error is extremely unlikely. The random 17-nucleotide portion of the BC is expected to be unique to each lentiviral integration event and thus tags all cells in a single clonal expansion. Note that the length of the barcode ensures a high theoretical potential diversity (∼4^17^ > 10^10^ barcodes per vector), so while the actual diversity of each Lenti-sgRNA/PuroR vector is dictated by the number of colonies generated during the plasmid barcoding step, it is very unlikely that we will observe the same BC in multiple clonal expansions.

FASTQ files were parsed using regular expressions to identify the sgID and BC for each read. To minimize the effects of sequencing error on BC identification, we required the forward and reverse reads to agree completely within the 23-nucleotide sequence to be further processed. Due to PCR and sequencing errors, genuine tumor barcodes can occasionally appear mutated. These “fake” tumor colonies are expected to be rare and sporadic. To minimize their impact, each barcode was converted into a 2D array of characters (n_barcodes x n_barcodes x barcode_length), and NumPy broadcasting was used to compute all pairwise Hamming distances simultaneously, while masking redundant comparisons. For barcode pairs with distances <=2, we retained the barcode with the higher read count.

After filtering, absolute cancer cell numbers for each barcode were calculated based on the regression of spike-in read counts against their known expected cell numbers. If not all expected spike-ins were detected in the sequencing data, missing spike-ins were assigned using the average ratio of observed counts to expected cell numbers across detected spike-ins. For samples with multiple spike-ins detected, linear regression without intercept was performed using the spike-in read counts and their expected cell numbers to calculate a conversion factor based on the slope of the regression line. Spike-ins with read counts below 10, relative counts outside ±50% of expected values, or high residuals (when R² < 0.9) were excluded to improve regression quality. This conversion factor was applied to all barcodes to estimate cell numbers, and barcodes with estimated cell numbers ≤1 were excluded from downstream analysis. For pre-transplantation samples lacking spike-ins, a default conversion factor of 10 cells per read was applied. The median sequencing depth across experiments was ∼1.3 reads per cell.

### Summary statistics for metastasis-related statistics

To assess the extent to which a given gene (*X*) *a*ffects metastatic clonal expansion, we compared the distribution of tumor sizes produced by vectors targeting that gene (sg*X* tumors) to the distribution produced by our negative control vectors (sg*Inert* tumors). We relied on two statistics to characterize these distributions: the size of tumors at defined percentiles of the distribution (specifically the 50^th^, 60^th^, 70^th^, 80^th^, 90^th^, and 95^th^ percentile tumor sizes), and the peak mode. The percentile sizes are nonparametric summary statistics of the tumor size distribution. The peak mode identifies the most common tumor size by determining the maximum of the kernel density estimate of log10-transformed cell numbers, computed using 512 evaluation points across the data range. This metric allows for quantification of the expansion of tumor colonies.

To quantify the extent to which each gene suppressed or promoted tumor growth, we normalized statistics calculated on tumors of each genotype to the corresponding inert statistic. The resulting ratios reflect the growth advantage (or disadvantage) associated with each tumor genotype relative to the growth of *sgInert* tumors.

For example, the relative i^th^ percentile size for tumors of genotype X was calculated as:

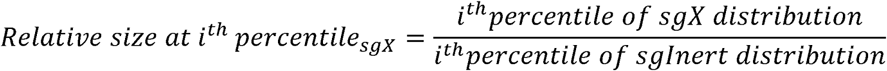

Likewise, the relative peak mode size for tumors of genotype X was calculated as:

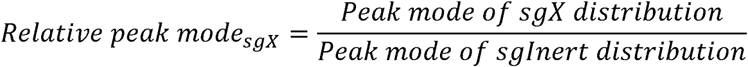

In addition to the tumor size metrics described above, we characterized the effects of gene inactivation on tumorigenesis in terms of the number of seeded tumors (“metastatic seeding”) and total neoplastic cell number (“metastatic burden”) associated with each genotype. Unlike the aforementioned metrics of tumor size, metastatic seeding and burden are linearly affected by pre-transplantation cell number and are thus sensitive to underlying differences in the representation of each barcoded tumor genotype in the pre-transplantation cells. Therefore, to assess the extent to which a given gene (*X*) affects metastatic seeding, we therefore first normalized the number of sg*X* tumors to the number of pre-transplantation sg*X* cells to account for differences in initial tumor transplantation:

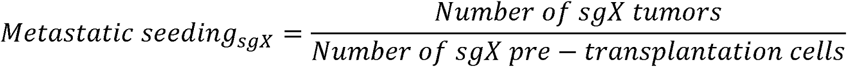

As with the tumor size metrics, we then calculated a relative metastatic seeding by normalizing this statistic to the corresponding statistic calculated using sg*Inert* tumors:

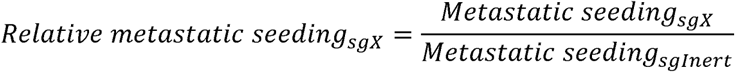

Genes that influence relative metastatic seeding modify the probability of tumor initiation and/or the very early stages of oncogene-driven epithelial expansion, which prior work suggests are imperfectly correlated with tumor growth at later stages. Relative metastatic seeding thus captures an additional and potentially important aspect of tumor suppressor gene function.

Analogous to the calculation of relative metastatic seeding, we characterized the effect of each gene on metastatic burden by first normalizing the sg*X* tumor burden to tumors to the number of pre-transplantation sg*X* cells:

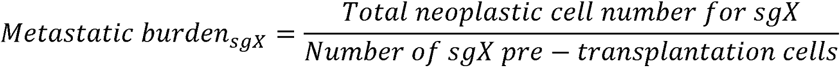

We then calculated a relative metastatic burden by normalizing this number to the corresponding statistic calculated using sgInert tumors:

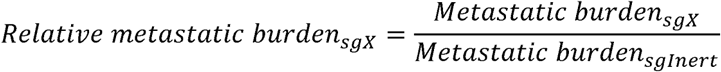

Metastatic burden is an integration over tumor size and number and thus reflects the total neoplastic load in each mouse. Metastatic burden is thus more strongly related to morbidity than are our metrics of tumor size and is closely related to traditional measurements of tumor progression such as duration of survival and tumor area. While intuitively appealing, tumor burden is notably noisier than our metrics of tumor size as it is strongly determined by the size of the largest tumors. To study the potential for aggressive metastatic colonies to release circulating tumor cells into the blood, we defined super-metastasis (“supermet”) as the percentage of disseminating tumors, defined as those tumor barcodes that were detected in both the target tissue and blood at matched time points within the same mouse, of all colonies:

In addition, we also calculated the peak mode of disseminating tumors to characterize the size at which tumors release cancer cells. As with the tumor size metrics, we normalized the supermet percentage calculated on tumors of each genotype to the corresponding inert statistic to reflect the impact on dissemination potential of each genotype relative to that of sg*Inert* tumors:

Finally, we also characterized the effects of gene inactivation on tumor arrest in the dormant phase b calculating the percentage of tumors that were found to have remained at a size considered to be dormant. As each metastatic colony expands from a single disseminated cell, we expected slow-growing dormant-like colonies to follow similar early growth trajectories independent of tissue or genotype. Therefore, we pooled tumor colonies across all tissues at endpoint, and calculated the size cutoff for tumors to be considered dormant by applying Gaussian mixture modeling to the distribution of log2-transformed tumor cell numbers of this pool. We fit a two-component GMM model with unequal variances (modelNames=”V”) using the mclust package, allowing us to establish a pooled cell number mode of dormant-like colonies and a valley point that represented the separator between dormant and expanding colonies. We then employed a constrained fitting approach using two-component GMM models on each specific tissue, genotype, and time point, where the mean of the dormant component was fixed to the pooled dormant mode and the point between dormant and expanding colonies was set to the pooled valley mode. This constraint improves model stability and ensures consistent identification of the dormant population across time. The constrained model uses an expectation-maximization (EM) algorithm with early stopping (convergence threshold: change in dormant proportion < 1×10^-6^ or maximum 50 iterations) to estimate the dormant and expanding components. The dormancy percentage statistic wa calculated as the percentage of dormant tumors:

We then calculated a relative dormancy percentage by normalizing this number to the corresponding dormancy percentage calculated using sg*Inert* tumors:

Furthermore, the relative statistics were converted to log2 fold-change for downstream analysis and visualization.

### Calculation of confidence intervals and P-values for tumor growth and number metrics

Confidence intervals and *P*-values were calculated using bootstrap resampling of tumors to estimate the sampling distribution of metastatic seeding, metastatic burden, and percentile sizes. 10,000 bootstrap samples were drawn for all reported P-values. 95% confidence intervals were calculated using the 2.5^th^ and 97.5^th^ percentile of the bootstrapped statistics. Because we calculate metrics of tumor growth that are normalized to the same metrics in sgInert tumors, under the null model where genotype does not affect tumor growth, the test statistic is equal to 1. Two-sided p-values were thus calculated as follows:

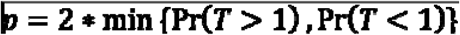

Where T is the test statistic and Pr(T>1) and Pr(T<1) were calculated empirically as the proportion of bootstrapped statistics that were more extreme than the baseline of 1. To account for multiple hypothesis testing, p-values were FDR-adjusted using the Benjamini-Hochberg procedure as implemented in th Python package statsmodels.

### Analysis of cBioPortal lung cancer genomic data

Clinical and genomic data were obtained from the cBioPortal for Cancer Genomics using the cBioPortalData R package. A combined cohort of 27 lung cancer studies was assembled, comprising 9,830 patients and 12,275 samples across multiple lung cancer subtypes. Mutation and copy number alteration (CNA) data for CXCR2 and RAC1 were downloaded from the cBioPortal query interface, and clinical data including overall survival (OS) status and duration and TNM staging were extracted for the combined cohort.

CXCR2 status was classified into three groups: knockout (KO), activating, and wild-type (WT). CXCR2 KO was defined as samples harboring homozygous deep deletions (GISTIC value = −2) or mutations at functionally critical residues, including the putative chemokine binding sites (residues 55, 104, 107, 123, 126-127, 196-201, 208, 211-212, 267, 274, 277, 289, 293, 296-297, 300) and the C-terminal phosphorylation region (residues 347-353). CXCR2 activating status was defined as high-level amplification (GISTIC value = 2). RAC1 status was similarly classified into three groups: KO was defined as homozygous deep deletion, activating was defined as high-level amplification or known oncogenic point mutations (P29S, P29L, Q61R, A159V, G12V, G12R, P34R, Q61K), and all remaining profiled samples were classified as WT. Samples not profiled for either gene were excluded from the respective analyses to ensure the WT group comprised only confirmed wild-type samples.

For samples present in multiple studies, a single representative entry was retained per sample, prioritizing the entry with the most complete clinical annotation for overall survival and metastatic staging. Metastatic staging was categorized from the TNM M-stage field as M0 (no distant metastasis) or M1 (distant metastasis present). The distribution of metastatic staging (M0 vs M1) across genotype groups was visualized using stacked bar plots showing the percentage of patients in each M-stage category. Overall survival was analyzed using Kaplan-Meier curves with the survfit function from the survival R package, and survival differences between groups were assessed using log-rank tests. All statistical analyses were performed in R version 4.5.2, and visualizations were generated using the survminer and ggplot2 packages.

### Statistical Analysis

Sample or experiment sizes were estimated based on similar experiments previously performed in our laboratory, as well as in the literature. For the experiments in which two or more cell types are co-cultured, we used at least 3 samples per group for FACS analysis, at least 10 images were taken per group for image quantification. Each experiment was repeated at least three times. All values are presented as means ± SD, with individual data points also shown in the figure. Comparisons of parameters between two groups were made by two-tailed Student’s t-tests. The differences among several groups will be evaluated by one-way ANOVA with Tukey-Kramer post hoc evaluation. P-values less than 0.05 and 0.01 were considered significant (*) or very significant (**), respectively.

All analyses of barcode sequencing data were performed in Python (3.10) and visualizations of data were performed in R (4.4.1). Sample sizes for Moba-seq experiments were determined based on previously reported power analyses^33^. Analyses of barcode sequencing data used non-parametric statistics; therefore, no assumptions about the distribution of data were made.

## Notes

### Competing Interest Statement

The authors have declared no competing interest.

### Summary of Updates

Figures 1, 5, and 6 revised and with new figure panels, supplemental figures updated

