## Supplementary figures and images for "CXCL-CXCR2 signaling drives cancer-endothelium interactions in SCLC metastatic seeding"

### Suppemental Figure 1

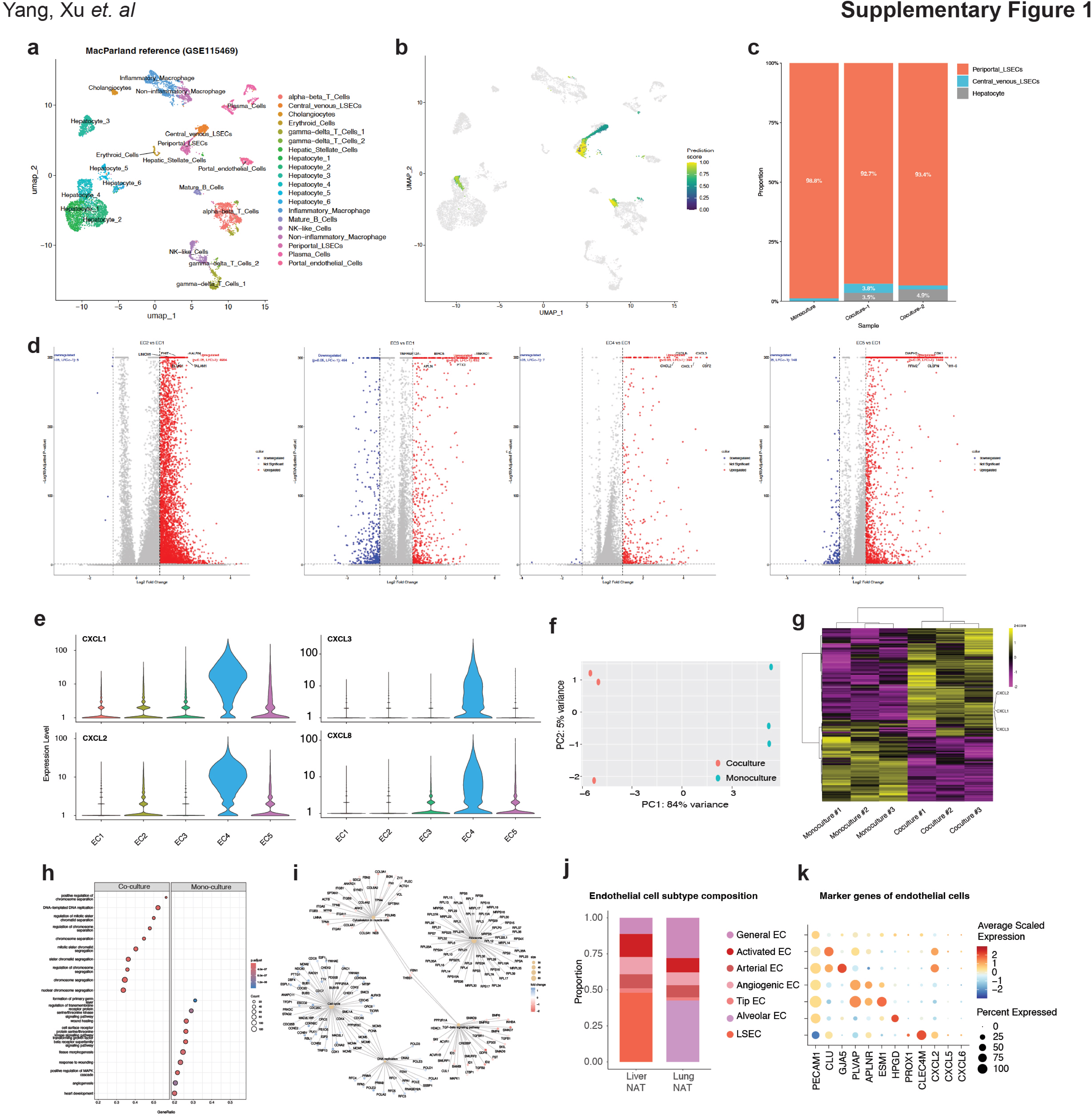

### Suppemental Figure 2

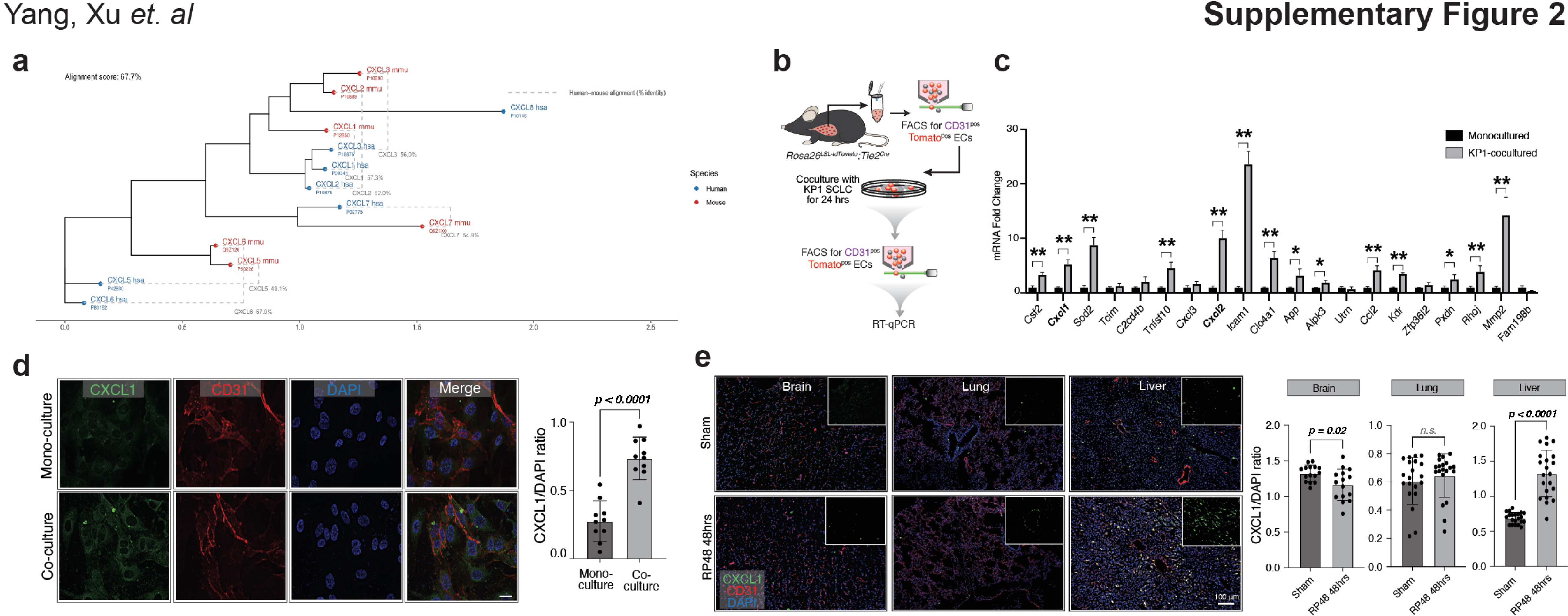

### Suppemental Figure 3

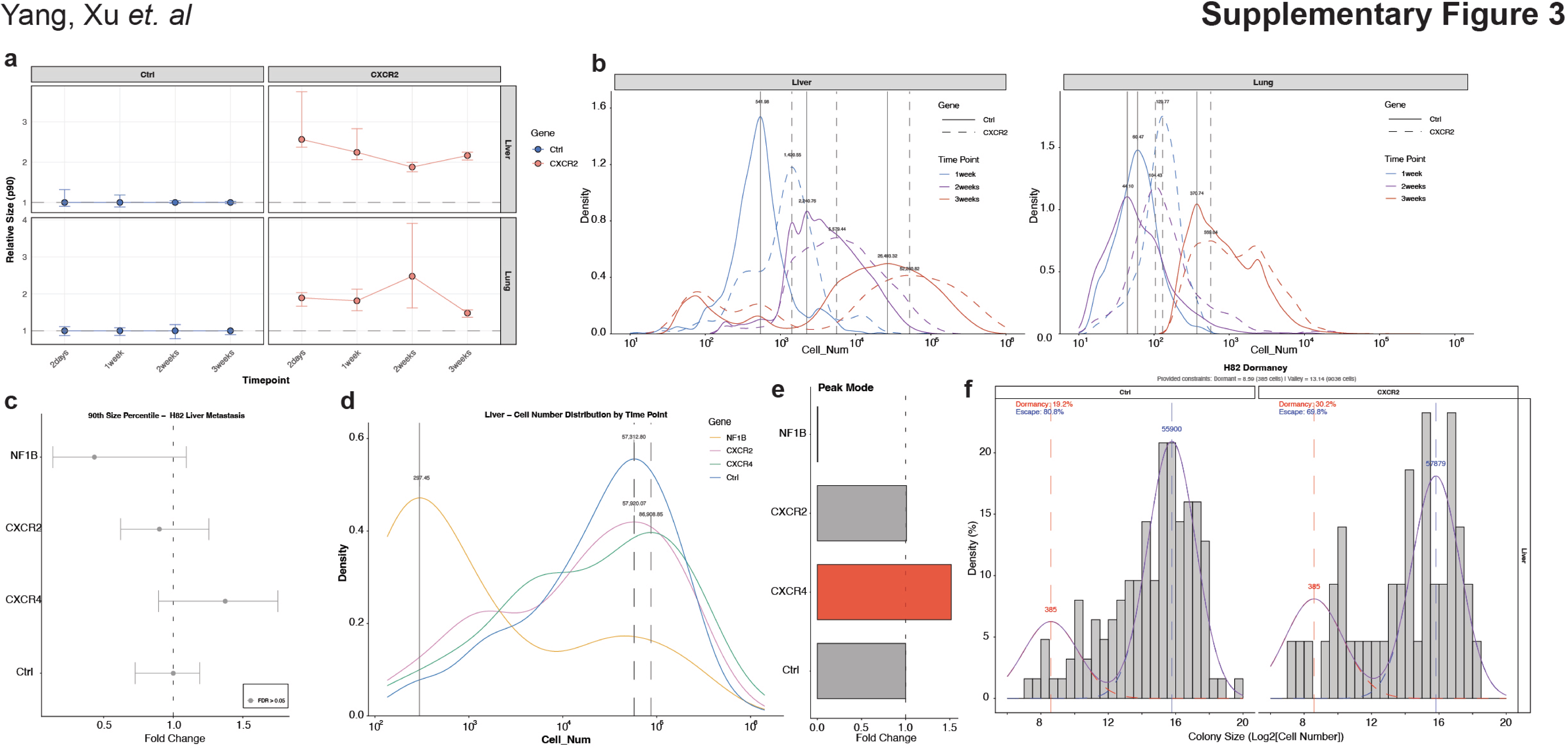

### Suppemental Figure 4

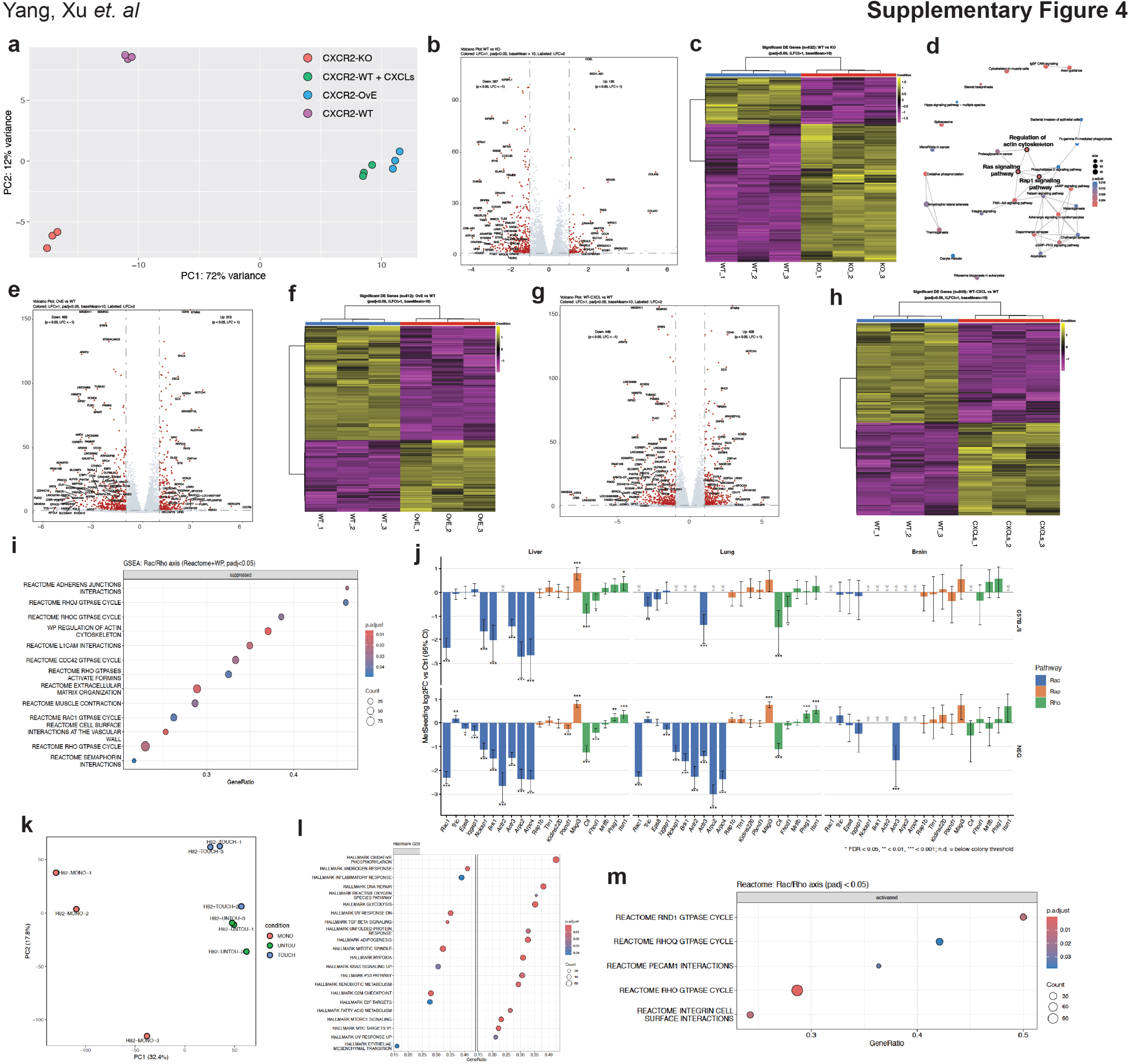

### Suppemental Figure 5

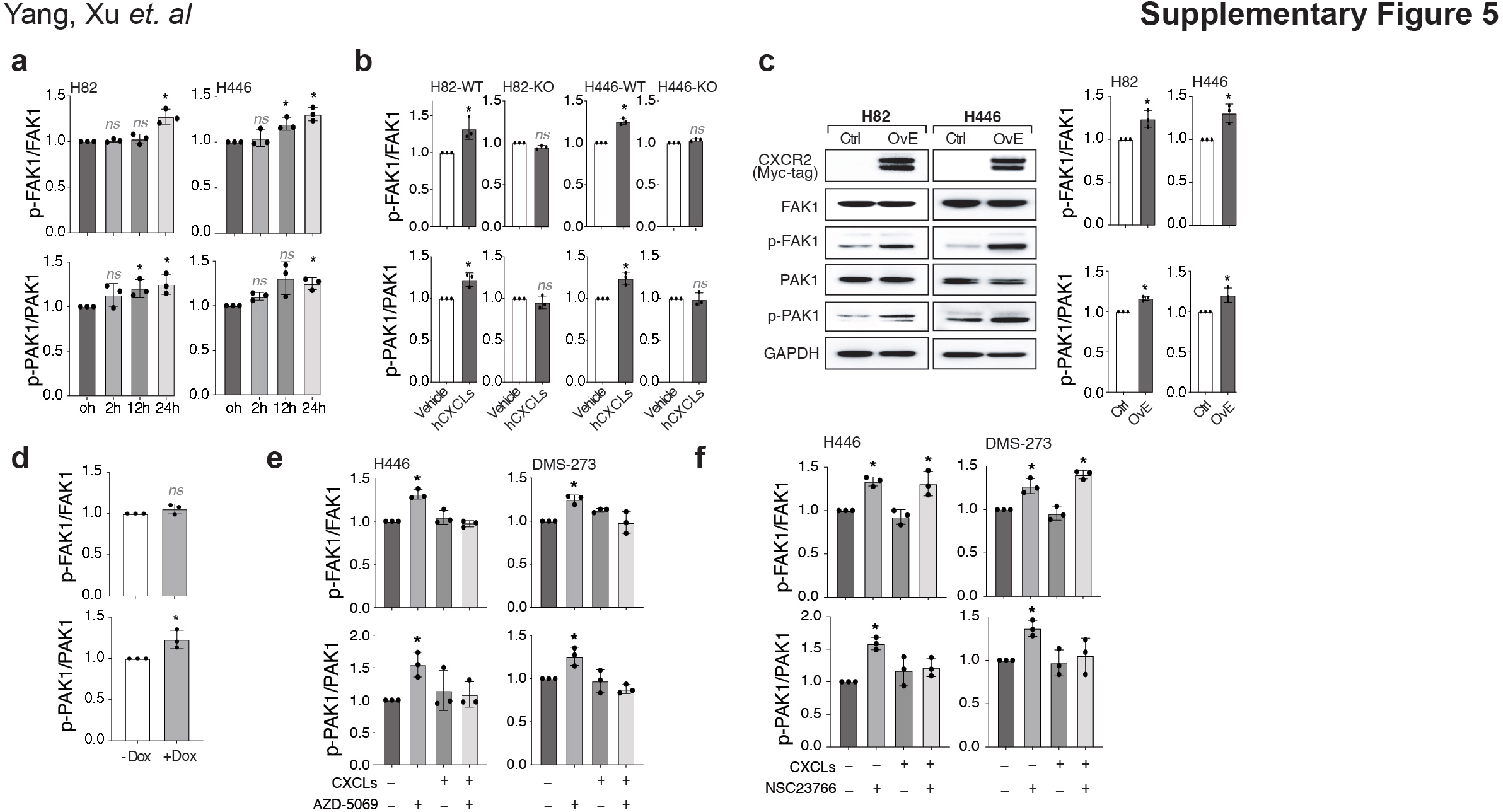

### Suppemental Figure 6

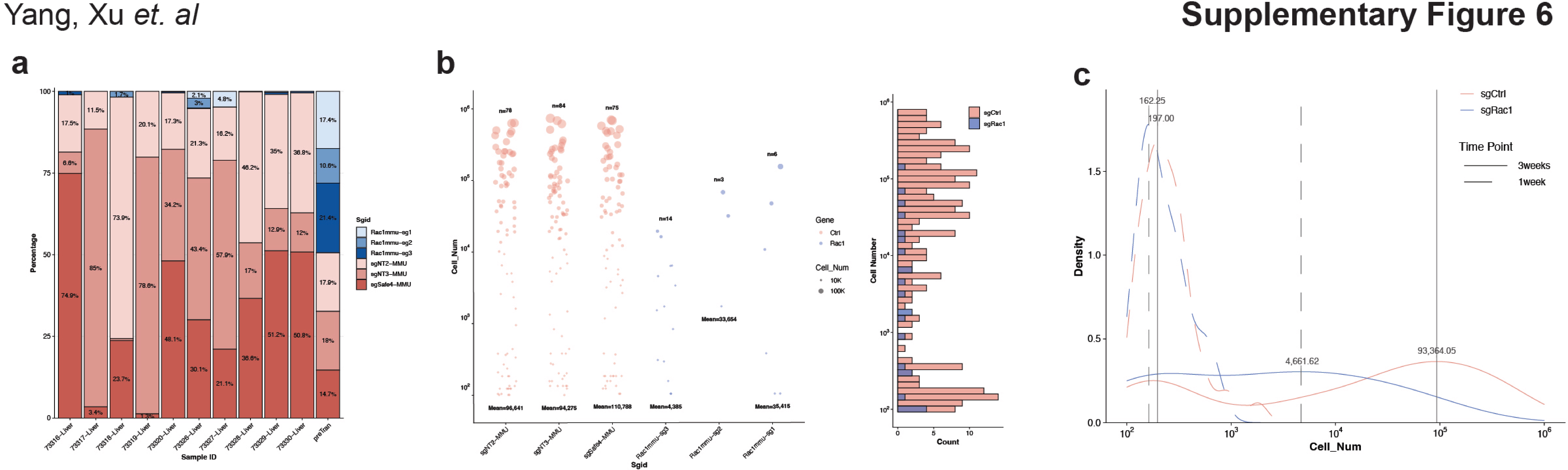

### Suppemental Figure 7

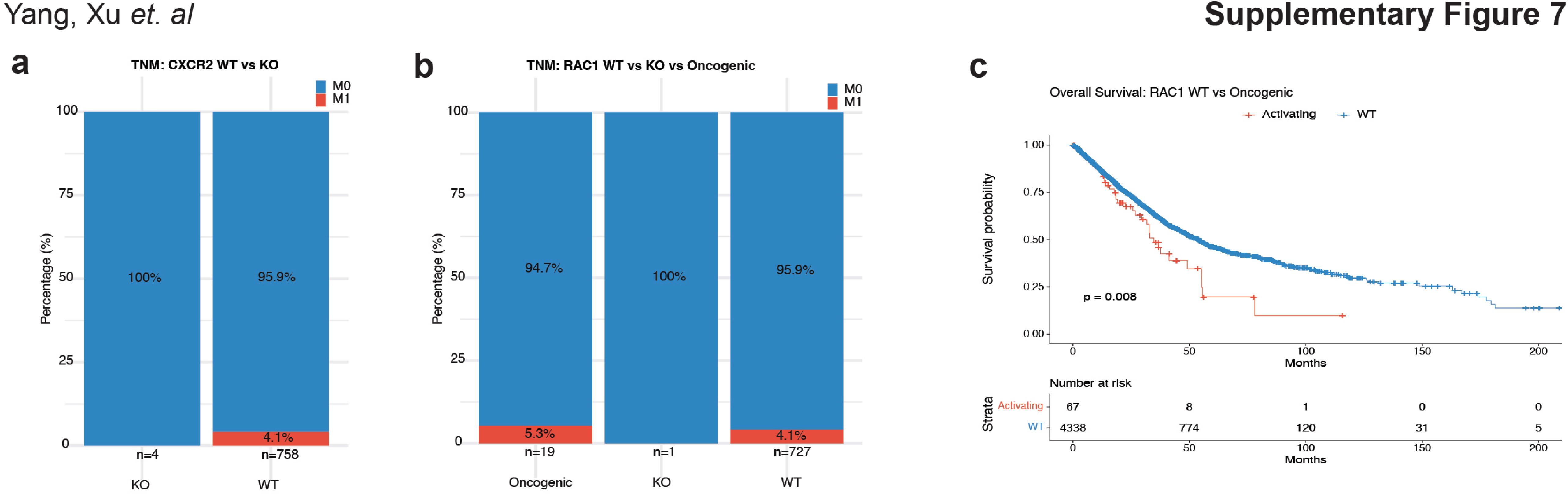
